# The staphylococcal type VII secretion system effector EsxC impacts daptomycin sensitivity through controlling bacterial cell envelope integrity

**DOI:** 10.1101/2023.11.20.567842

**Authors:** Victoria Smith, Giridhar Chandrasekharan, Robeena Farzand, Kate E. Watkins, Ramon Garcia Marset, Jeannifer Yap, Sebastien Perrier, Arnaud Kengmo Tchoupa, Meera Unnikrishnan

## Abstract

The human pathogen *Staphylococcus aureus* encodes a specialised type VII secretion system (T7SS), which plays an important role in bacterial virulence during infection. However, the functions the T7SS during infection and in bacterial physiology remain unclear. Here we demonstrate that *S. aureus* strains lacking the the T7SS effector EsxC (Δ*esxC*) was highly sensitive to the important last resort drug, daptomycin, as well as other membrane-targeting antibiotics, including gramicidin and bithionol. To understand how EsxC mediates increased antibiotic sensitivity, we investigated its functions in the staphylococcal cell envelope. Scanning electron microscopy analysis of an *esxC* mutant revealed a distinct cell surface morphology. Interestingly, Δ*esxC* displayed a decrease in membrane fluidity, altered membrane protein profiles and altered cell wall synthesis. The *esxC* mutant demonstrated enhanced daptomycin binding which correlated with the increased negative charge of mutant membranes. Calcium ions, which can bind membranes affecting charge, impacted growth of Δ*esxC* and sensitivity to daptomycin, suggesting that EsxC may modulate calcium binding to membranes. Furthermore, the *esxC* mutant displayed a heightened susceptibility to daptomycin during intracellular infection, and in a murine skin infection model. Thus, our data show that the T7SS effector EsxC impacts sensitivity of *S. aureus* to membrane-acting drugs such as daptomycin through modulation of cell membrane integrity, indicating its potential as a drug target.

**Author Summary:** T7SS has a range of functions in bacteria including specific roles in bacterial physiology including DNA uptake, membrane integrity and bacterial development. In *S. aureus* T7SS has been shown to be critical for bacterial virulence, intra-species competition and in host cell interactions, although their functions in bacterial physiology are not clear. Here we report a role of the staphylococcal T7SS effector EsxC in the modulation of the cell membrane and surface integrity, which impacts the activity of membrane targeting drugs like daptomycin. Our data indicate that targeting this system could potentially enhance activity of existing therapeutic agents.

## Introduction

*S. aureus* is a highly versatile pathogen and a commensal bacterium which causes infections in both humans and animals. Staphylococcal infections can be nosocomial or community-acquired, and can range from mild skin abscesses to severe life-threatening infections including septic shock and bacteraemia (1). *S. aureus* infections place a huge cost burden on healthcare sectors across the world, especially due to the occurrence of antibiotic-resistant strains, such as methicillin-resistant *S. aureus* (MRSA) and strains with varying levels of vancomycin resistance (2, 3).

*S. aureus* possesses an array of surface and secreted factors that help the bacterium colonise the host and establish an infection (4). One such virulence-associated factor is the type VII secretion system (T7SS), which is widely found in Gram positive bacteria (5). The T7SSb in *S. aureus* and other firmicutes is distinct from the systems in mycobacteria (T7SSa) although both are characterised by small substrates containing the WXG motif, such as EsxA and EsxB (6, 7). The T7SS in *S. aureus* is modular in nature, with heterogenous expression between strains (8, 9). Typically, in commonly studied *S. aureus* strains including USA300, Newman and RN6390, the T7SS locus comprises of integral membrane proteins (EsaA, EssA, EssB and EssC), cytosolic proteins (EsaB and EsaG), several secreted substrates (EsxA, EsxB, EsxC, EsxD and EsaD) and chaperones (EsaE) (7, 10). EssC is the central transporter which is responsible for export of substrates and co-dependent export of secreted substrates has been reported (6, 11). While specific interactions between the different T7SS components have been described, the overall structure of the secretion apparatus is poorly defined for the *S. aureus* T7SS (10, 11). Assembly of a functional system is dependent on the membrane microdomain protein, flotillin A (FloA)(12).

Recent studies have reported a role for T7SS in modulation of intraspecies competition. A T7SS substrate EsaD with nuclease activity mediates killing of *S. aureus* strains that lack the anti-toxin EsaG (13). An LXG-domain containing protein, TspA, encoded distally to the T7SS cluster shows T7SS-dependent toxic activity, which can be neutralized by an antitoxin TsaI (14). Some substrates modulate host interactions during infection; EsxA modulates host cell death and EsaE elicits cytokine production during infection (15, 16). However, effectors of this system are yet to be defined in many *S. aureus* strains and their specific functions remain elusive.

T7SS in other bacterial species has been described to support bacterial functions like DNA transfer, membrane integrity and spore development, although for *S. aureus*, it is not clear if T7SS is important in staphylococcal physiology (17–19). Our recent study on the T7SS substrate EsxC showed that a mutant lacking this protein was more sensitive to unsaturated host-derived fatty acids (20). T7SS defects compromised membrane integrity and induced oxidative stress in presence of antimicrobial fatty acids, indicating a role for these proteins in membrane homeostasis. Here we report that the absence of EsxC causes an increased sensitivity to membrane targeting drugs like daptomycin, both during infection *in vitro* and *in vivo*. We demonstrate distinct effects on cell surface morphology, membrane fluidity and charge, indicating a role for T7SS in controlling membrane integrity potentially through modulating calcium binding. Our data suggest that targeting the T7SS could augment activities of membrane-acting drugs like daptomycin.

## Results

### A mutant lacking T7SS effector EsxC is more sensitive to membrane-acting drugs

The cyclic lipopeptide antibiotic, daptomycin, is a key drug used to treat resilient *S. aureus* infections. Daptomycin binds to the cell membrane in a calcium-dependent manner (21). As our previous studies indicated a potential impact of T7SS, in particular the effector EsxC, on membrane homeostasis (20), we investigated whether the *S. aureus* USA300 JE2 mutants lacking the T7SS substrate EsxC (Δ*esxC)* respond differently to membrane-acting drugs like daptomycin. Growth of Δ*esxC* was compared to the WT USA300 JE2 strain in the presence or absence of daptomycin. When cultured in presence of 5 μg/ml daptomycin, growth of the WT USA300 JE2 strain was delayed, but growth of USA300 JE2 Δ*esxC* was significantly impaired compared to the WT (Fig. 1 A). No differences in growth rates were seen between the strains in tryptic soy broth (TSB) without antibiotic. Further, a 10-fold decrease in colony forming unit counts for the mutant was also observed in the presence of daptomycin (Fig. 1 B). Complementing Δ*esxC* with pOS1-*esxC* restored WT phenotype (Fig. 1 C, D), although daptomycin did not inhibit the pOS1 empty plasmid-bearing Δ*esxC* to the same extent as Δ*esxC* without plasmid. This could be attributed to differences in growth profiles of pOS1 containing WT and mutant strains, possibly due to the effects of chloramphenicol exposure prior to performing growth assays. Next, a daptomycin killing assay was employed where logarithmic phase bacteria were treated for 2 h with daptomycin. Δ*esxC* was more sensitive to daptomycin in comparison to the WT in this killing assay, with the Δ*esxC* pOS1-*esxC* strain reversing daptomycin sensitivity (Fig. 1 E, F). When we stained bacteria with propidium iodide (PI), a fluorescent intercalating agent which cannot pass through the membranes of viable cells, cytosols of daptomycin-treated Δ*esxC* showed a higher intensity of PI compared to the WT, as quantified by fluorescence microscopy (Fig. 2 A, B). Staining with the lipid staining dye FM1-43 was of similar intensity for WT and Δ*esxC*, although it appeared less ordered in the daptomycin-treated Δ*esxC* sample (Fig. 2 A, B). Thus, Δ*esxC* demonstrated increased membrane permeability in the presence of daptomycin.

**Figure 1.**
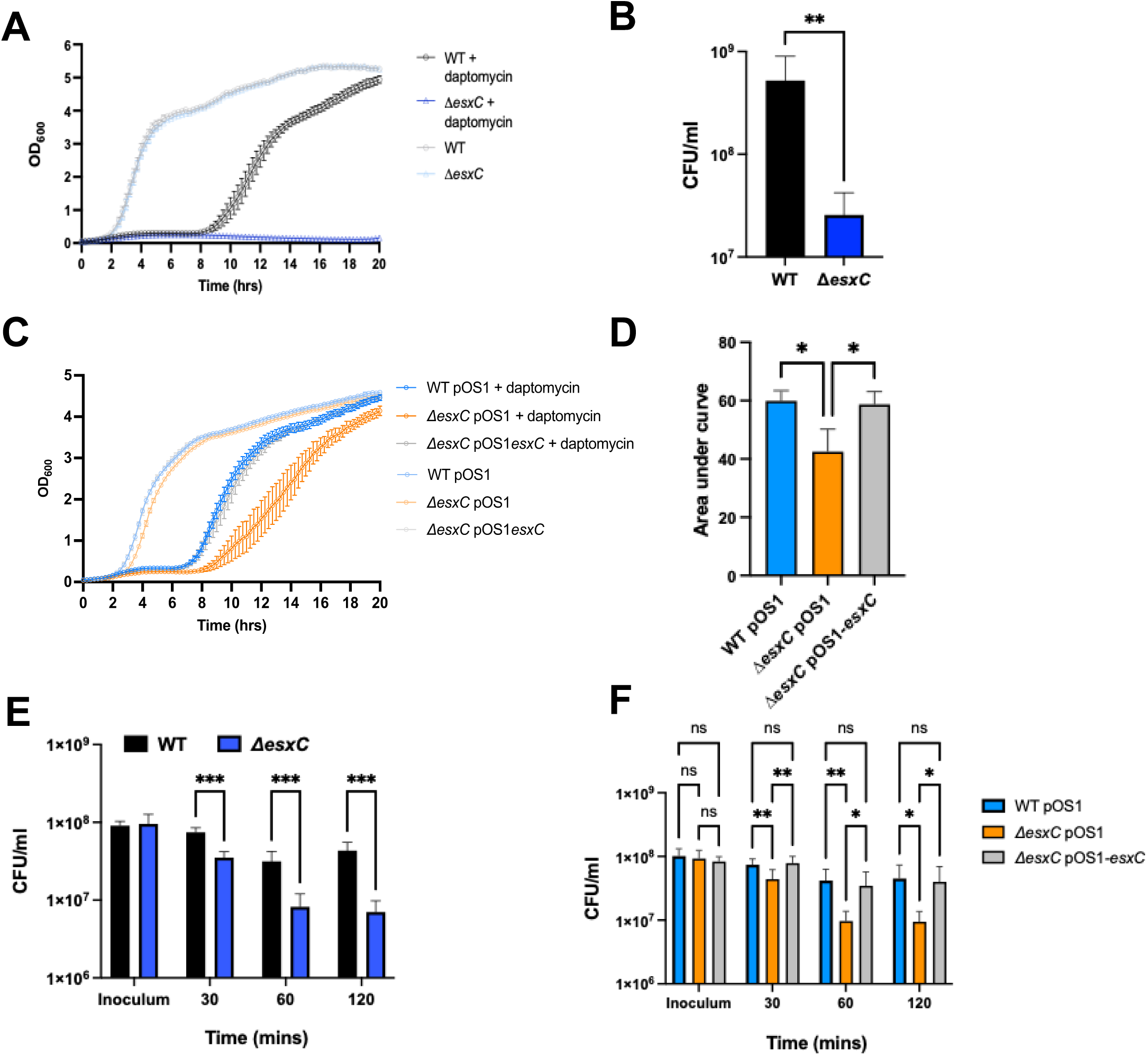
An *esxC* mutant is highly sensitive to daptomycin. **A)** Growth curves of *S. aureus* WT USA300 JE2 and Δ*esxC* in TSB in the absence or presence of 5 μg/ml daptomycin and 1 mM CaCl_2_. Mean ± standard error of the mean (SEM) is shown, N = 3 (biological replicates). **B)** CFU/ml was enumerated after 8 h daptomycin treatment. N = 3 (biological replicates), mean + standard deviation (SD), *** P ≤ 0.001 using an unpaired t-test **C)** Growth curves of USA300 JE2 WTpOS1, Δ*esxC*pOS1 and complemented strain, Δ*esxC*pOS1-*esxC* in TSB supplemented with 5 μg/ml daptomycin and 1 mM CaCl_2_. Mean ± SEM shown, N = 3 (biological replicates); AUC was calculated for the strains grown in presence of daptomycin **(D)** Mean + SD ** P ≤ 0.05 using a one-way ANOVA with Tukey’s multiple comparison test. **E)** CFU/ml of WT USA300 JE2 and Δ*esxC* were determined from a killing assay in the presence of 10 μg/ml daptomycin and 1 mM CaCl_2_ at different timepoints. **F)** CFU/ml from killing assay using WT USA300 JE2, Δ*esxC* and Δ*esxC* complemented strains in the absence or presence of 10 μg/ml daptomycin and 1 mM CaCl_2_ at different timepoints, mean ± SD, N = 3 (biological replicates), * P ≤ 0.05, ** P ≤ 0.01, *** P ≤ 0.001 using a two-way ANOVA with Tukey multiple comparison test.

**Figure 2.**
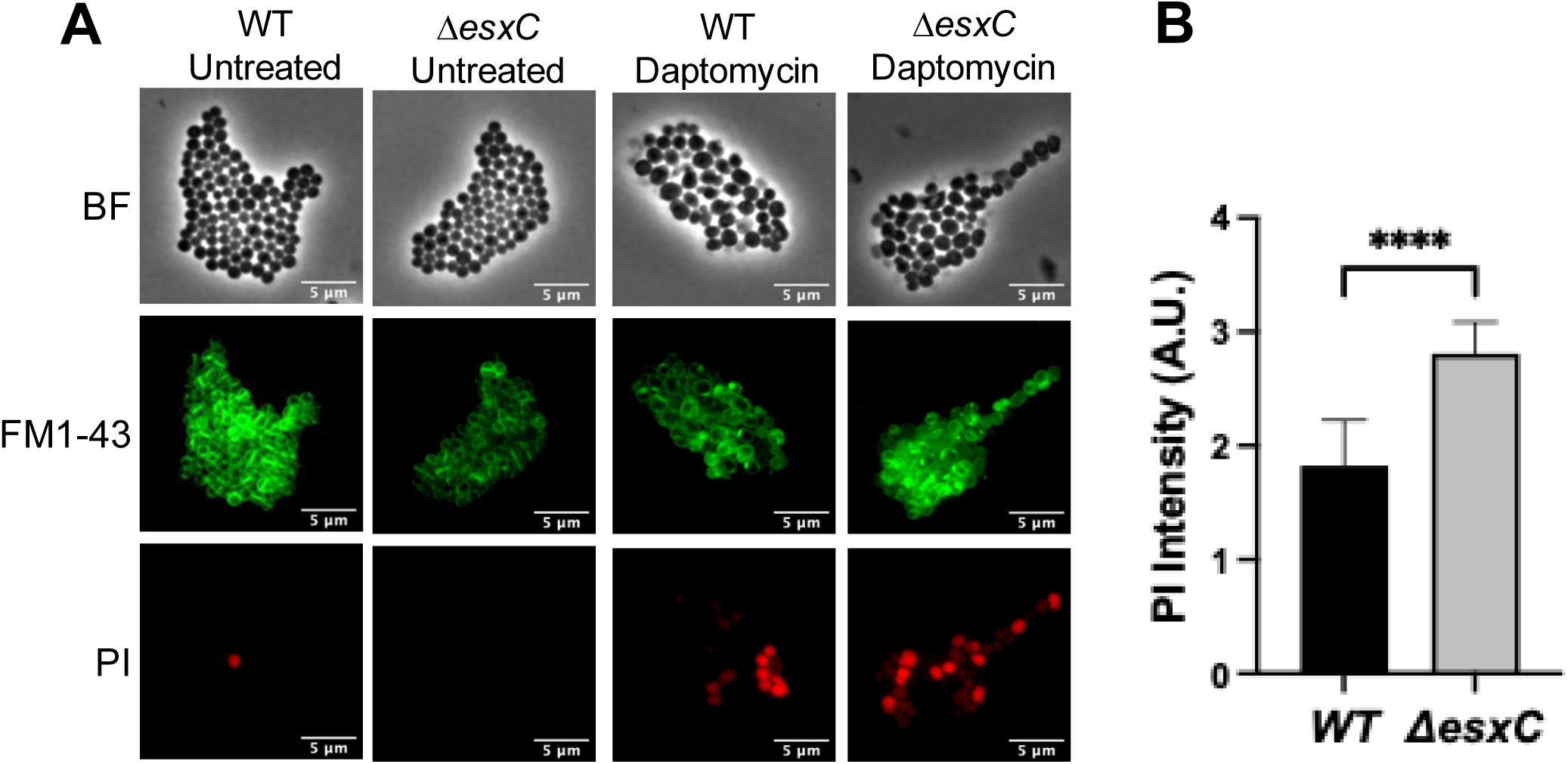
Membrane permeability of the *esxC* mutant in the presence of daptomycin. **A)** Representative fluorescent micrographs of *S. aureus* USA300 JE2 WT and Δ*esxC* treated with daptomycin. Cell membranes were stained by FM1-43 and imaged using FITC filter channel, PI was imaged using Texas red filter channel. **B)** Images of daptomycin-treated samples (A) were quantified using FIJI. Mean fluorescence intensity was quantified from 10 images from each experiment and normalised to the number of cells in the field of view. Data presented is from 3 independent experiments. ****P < 0.0001 using an unpaired t test.

To investigate whether other T7SS effectors had similar effects, we studied strains lacking substrates EsxA and EsxB. We used mutants in USA300 LAC and Newman backgrounds, as USA300 JE2 mutants in these genes were not available. A defect in growth was seen in USA300 LAC Δ*esxA* mutant compared to the USA300 LAC WT strain, while Δ*esxB* did not appear to have a growth defect (Fig. S1 A, B). The Newman strain was more sensitive than USA300 JE2 to daptomycin, with growth being delayed until ∼14 h. Again a growth defect was observed for the Δ*esxA* mutant, but not the Δ*esxB* mutant in presence of daptomycin (Fig. S1 C, D). Furthermore, a USA300 JE2 mutant lacking another T7SS substrate, EsxD did not display any growth defects in the presence of daptomycin (Fig. S1 E,F). We also studied a strain lacking and the central transporter EssC (*ΔessC*) responsible for transport of T7SS effectors. As expected, the deletion of *essC* significantly impaired the growth phenotype compared to the WT (Fig S1 G,H). We further studied the effect of EsxC on daptomycin sensitivity in other strain backgrounds. The growth of RN6390 *S. aureus* was impacted more by the addition of daptomycin, with the bacterial growth initiating much later (∼14 h) compared to the USA300 JE2 strain (∼8 h). However, similar to the JE2 mutants, RN6390 Δ*esxC* and *ΔessC* mutants were more susceptible than the WT to daptomycin (Fig. S1 I, J).

To determine if the differences observed between the WT and Δ*esxC* were limited to daptomycin, other membrane-targeting antimicrobials, such as gramicidin and bithionol, were tested. Gramicidin is a polypeptide antimicrobial peptide that forms ion channels in the cell membrane allowing diffusion of monovalent ions such as potassium (K^+^) and sodium (Na^+^), disrupting cell ion concentrations. Bithionol is an inhibitor of soluble adenylyl cyclase and its membrane-disrupting activity was associated with its ability to kill *S. aureus* persister cells (22). When treated with 2 μg/ml gramicidin, the WT JE2 strain did not have any growth defects, however the Δ*esxC* mutant displayed slower growth (Fig. 3 A, B). A similar defect in growth was also seen in the presence of biothionol, although only a slightly slower growth was seen for the mutant in the presence of 1 μg/ml bithionol (Fig. 3 C,D). Finally, this effect appears to be specific to membrane-acting drugs as other drugs which target the cell wall (vancomycin, oxacillin) or DNA replication (ciprofloxacin, mitomycin) did not differentially affect the growth of the Δ*esxC* (or Δ*essC*) as compared to the WT (Fig. S2).

**Figure 3.**
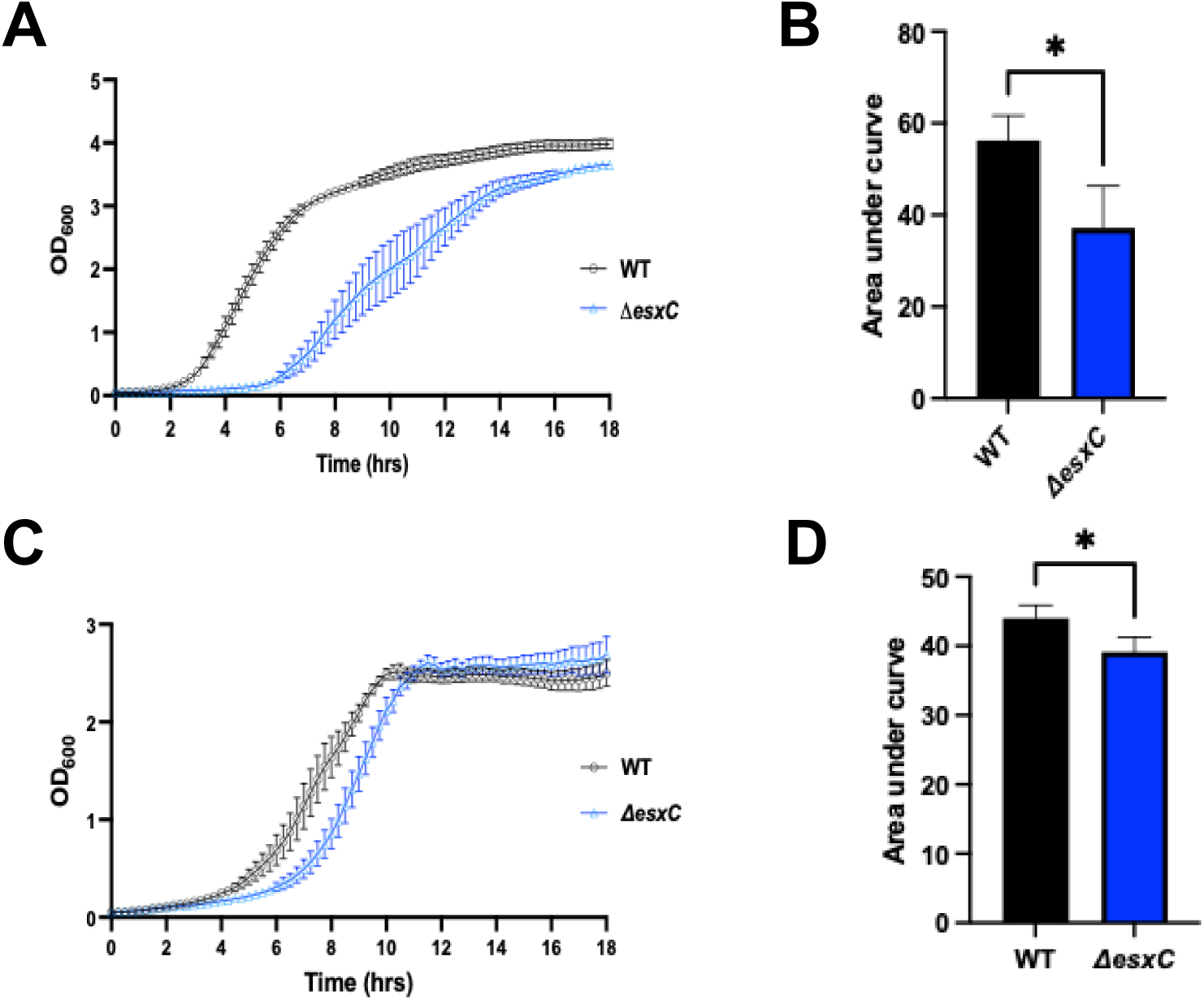
The *esxC* mutant show higher sensitivity to other membrane acting drugs. **A)** Growth curves of USA300 JE2 WT and Δ*esxC* in the presence of 2 μg/ml gramicidin, Mean ± SEM, N = 3 (biological replicates). **B)** Area under curves (AUC) were calculated and the mean AUC + SD was plotted for growth curves in (A). **C)** Growth curves of USA300 JE2 WT and Δ*esxC* in the presence of 1 μg/ml bithionol. Mean ± SEM, N = 3 (biological replicates). **D)** AUC were calculated and the mean AUC + SD was plotted for growth curves in (C). *P ≤ 0.05 using an unpaired t-test.

### *ΔesxC* displays decreased membrane fluidity and altered membrane protein profiles

Changes in membrane fluidity can affect membrane function and antibiotic sensitivity. To study the effect of EsxC on membrane fluidity, we assessed the membrane fluidity of the WT and Δ*esxC* mutant using multiple assays. WT and Δ*esxC* were stained with a fluorescent membrane dye DiI12C, which has been used detect regions of increased fluidity in bacteria (23). As reported previously for *S. aureus*, we observe a few fluid regions in the WT, and an uneven staining of the membrane. In contrast, a very weak staining of the membranes was observed for the Δ*esxC* mutant (Fig. 4 A), which may indicate a decrease in membrane fluidity. An alternate assay using pyrene decanoic acid, an excimer-forming lipid (24), was employed to measure the fluidity of WT and Δ*esxC* membranes. Compared to the WT, the cytosolic membrane of the Δ*esxC* mutant showed a mild but statistically significant increase in rigidity (Fig. 4 B). Finally, membrane fluidity was measured using the fluorescent dye Laurdan (6-dodecanoyl-2-dimethylaminonaphthalene) which is incorporated into the membrane bilayer (25). Logarithmic phase WT cells treated with laurdan showed a lower average generalized polarisation value of 0.33 compared to 0.42 for *ΔesxC*, which was reversed in the complemented strain Δ*esxC* pOS1*esxC*, suggesting that the membrane of the *ΔesxC* mutant is more rigid in comparison to that of the WT (Fig. 4 C). Overall, the data indicate that EsxC contributes to modulation of *S. aureus* membrane fluidity.

**Figure 4.**
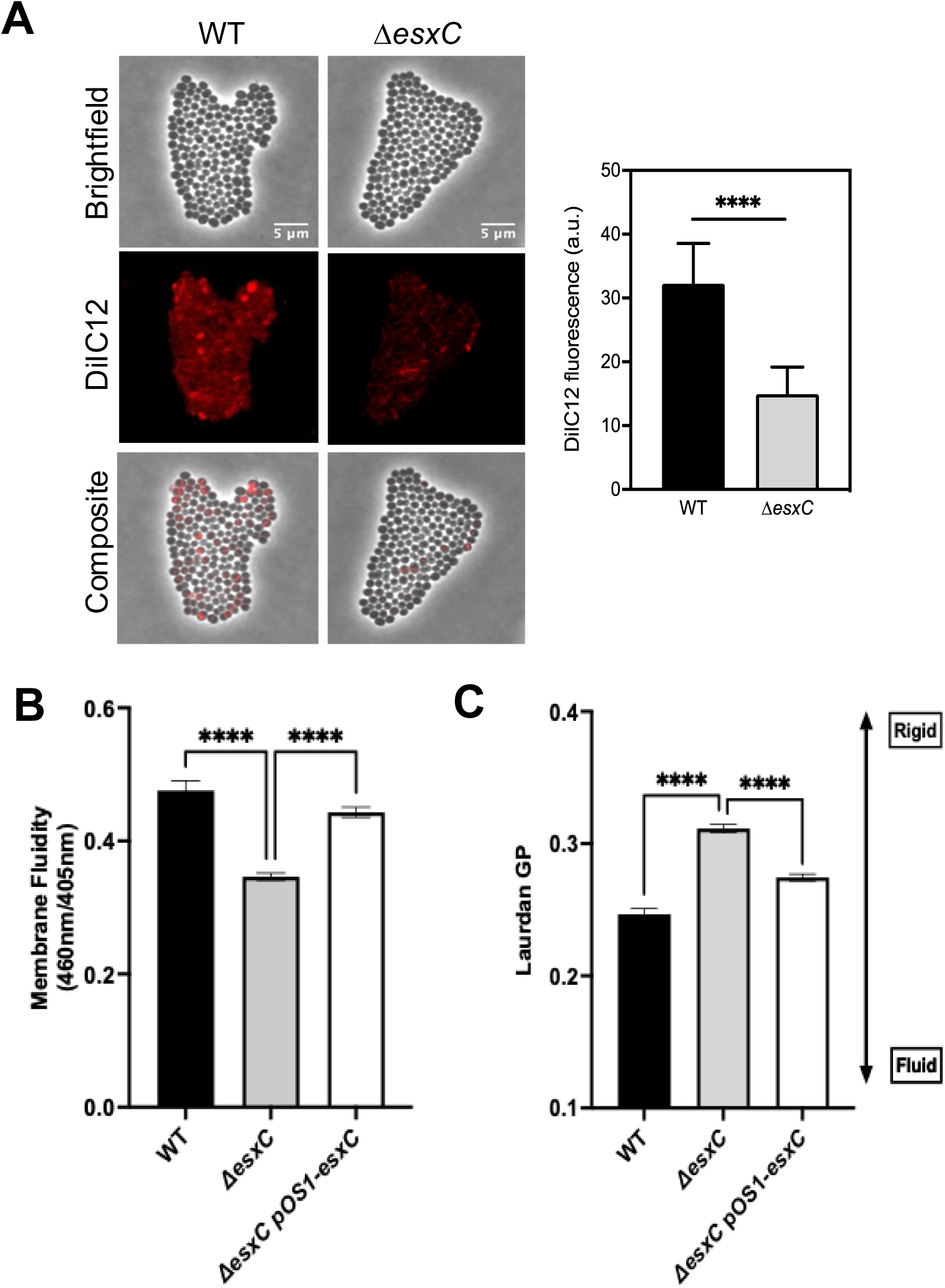
EsxC affects membrane fluidity. **A)** Widefield micrographs of *S. aureus* WT USA300 JE2 and Δ*esxC* after growth in TSB to OD_600_ of 1.0 in the presence of the lipophilic dye DiIC12. Images are representative of 4 independent experiments. The DiIC12 fluorescence of 80 bacterial clusters from different fields per strain was quantitated with ImageJ. Means + SD are shown, N = 4 (biological replicates); **** P < 0.0001, using an unpaired t-test. **B)** The membrane fluidity of *S. aureus* WT USA300 JE2, Δ*esxC* and Δ*esxC* pOS1-*esxC* as measured with a pyrene decanoic acid staining-based assay. Mean ± SEM are shown, N = 6 (biological replicates), * indicates P < 0.05 using a one-way ANOVA. **C)** USA300 JE2 WT, Δ*esxC* and Δ*esxC* pOS1-*esxC* were grown to logarithmic phase in TSB at 37°C prior to staining with Laurdan. To assess membrane fluidity, Laurdan generalised polarisation (GP) was calculated using the equation (I_460_ – I_500_)/ (I_460_ + I_500_) where I refers to the fluorescence intensity at the indicated emission wavelength. *** P <0.0001 using a one-way ANOVA, N = 3 (biological replicates), mean + SD.

Previously, we reported that the mutants Δ*esxC* and Δ*essC* had distinct total protein profiles with and without treatment with linoleic acid (20). Given the changes seen in bacterial membranes in the absence of EsxC, we examined global changes in the protein content in the membrane. Membrane fractions extracted from logarithmic phase (OD_600_ = 3.0) cultures of WT and Δ*esxC* were analysed by nanoLC-ESI-MS/MS. 13 proteins were altered in abundance [log2 (fold change) = 1.0, *P* < 0.05] in Δ*esxC* in relation to the WT (Table 1). Membrane proteins were increased in abundance and EssD, a T7SS substrate and nuclease, was decreased significantly in the membranes of Δ*esxC.* A secretome analysis from culture supernatants was also performed, which showed a lower abundance of substrates EsxA and EssD in Δ*esxC* compared to WT (Table S1). Thus, the *esxC* mutant show distinct changes to the overall membrane protein profiles and secretion of certain T7SS substrates appear to be co-dependent on substrate EsxC.

**Table 1.** Proteins significantly changed in Δ*esxC* mutant membrane preparations relative to WT USA300 JE2.

| Uniprot ID | $\log_2$ fold change | Adjusted $P$ value | Description |
| --- | --- | --- | --- |
| A0A0H2XJR2 | -5.46 | 6.21E-08 | Pyridine nucleotide-disulfide oxidoreductase |
| A0A0H2XJ15 | -5.34 | 3.80E-09 | TPR domain protein |
| A0A0H2XIV9 | -5.33 | 1.49E-07 | T7SS protein, EssD |
| A0A0H2XJ90 | -4.82 | 6.92E-09 | D-isomer specific 2-hydroxyacid dehydrogenase family protein |
| A0A0H2XF36 | 3.62 | 2.14E-08 | Uncharacterised protein |
| A0A0H2XJV3 | 3.67 | 2.26E-08 | ABC transporter, ATP-binding protein |
| A0A0H2XGS2 | 3.70 | 1.71E-08 | Putative sensor histidine kinase |
| A0A0H2XG87 | 4.01 | 1.59E-08 | Bifunctional ligase/repressor BirA |
| A0A0H2XH85 | 4.14 | 1.04E-08 | Cobalamin synthesis protein/P47K family protein |
| A0A0H2XER9 | 4.38 | 9.36E-09 | LysR family regulatory protein |
| A0A0H2XEB7 | 5.25 | 2.39E-07 | Integral membrane protein |
| A0A0H2XJ96 | 5.27 | 1.04E-08 | NADH-dependent flavin oxidoreductase |
| A0A0H2XHU5 | 5.97 | 2.55E-09 | Putative membrane protein |

### EsxC displays altered cell surface morphology and decreased rate of cell wall synthesis

As EsxC appeared to be involved in membrane function, we next examined the cell surface of Δ*esxC* using scanning electron microscopy. The cell surface of WT grown to early logarithmic phase have the typical smooth spherical shape of *S. aureus*. However, Δ*esxC* appears deflated with dents or hollows (Fig. 5 A). The WT phenotype was restored in Δ*esxC* pOS1-*esxC* complemented strain (Fig. 5 B). Interestingly, this cell surface defect is not seen in all T7SS mutants; strains lacking EsxD (Δ*esxD*) did not have a defective cell surface phenotype (Fig. S3). *S. aureus* cell surface alterations were seen in USA300 (LAC) and Newman strains lacking EsxA (*ΔesxA)*, but not in *ΔesxB* (Fig. S3). However, RN6390 strains lacking *esxC* showed very little surface defects (Fig. S3). Finally the JE2 transporter mutant Δ*essC* showed defects similar to Δ*esxC* (Fig S3). Differences seen in surface morphologies between WT and mutants were not due to growth rate differences, as growth rates of all mutant strains in the different strain backgrounds (*ΔessC*, *ΔesxC*, *ΔesxA*, Δ*esxB,* and *ΔesxD)* in TSB were similar to that of their WT (Fig. S1). Our data suggest that the cell envelopes of mutants lacking specific T7SS components were altered, potentially due to a defect in the cell membrane and/or cell wall structure.

**Figure 5.**
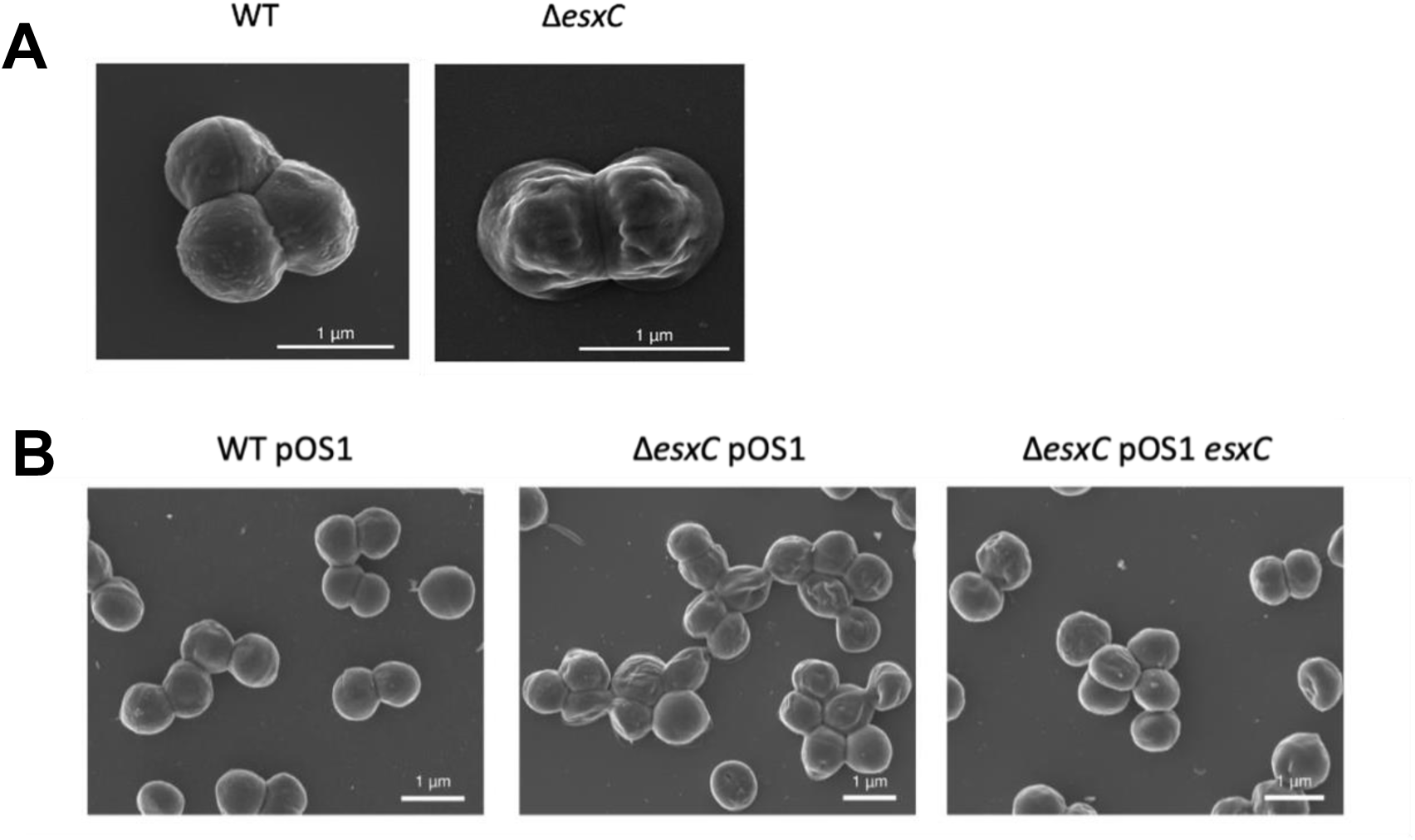
The *esxC* mutant displays distinct surface morphologies. **A)** Representative SEM images of *S. aureus* WT USA300 JE2 and *ΔesxC* grown to early logarithmic phase. **B)** Representative SEM images of *S. aureus* USA300 JE2 WT pOS1, *ΔesxC* pOS1, *ΔesxC* pOS1-*esxC* grown to early logarithmic phase.

To investigate if the surface structure changes were associated with a defect in cell wall structure or synthesis, transmission electron microscopy was undertaken. However, there were no clear differences in cell wall thickness between strains (Fig. S4). To study whether peptidoglycan synthesis and accumulation were affected in the T7SS mutants, HADA, a fluorescent D-amino acid was used (26). The dye was either added to a fresh subculture and grown until the bacteria reached logarithmic phase or added for 10 min once the culture reached an OD_600_ of 1. No differences were observed in the cells which had been grown with TSB supplemented with HADA (Fig. 6 A, B), however addition of HADA during exponential phase growth revealed a faster accumulation of HADA in the WT compared to *ΔesxC* (Fig. 6 A, C). This suggests that T7SS mutants displayed a decreased rate of new peptidoglycan synthesis.

**Figure 6.**
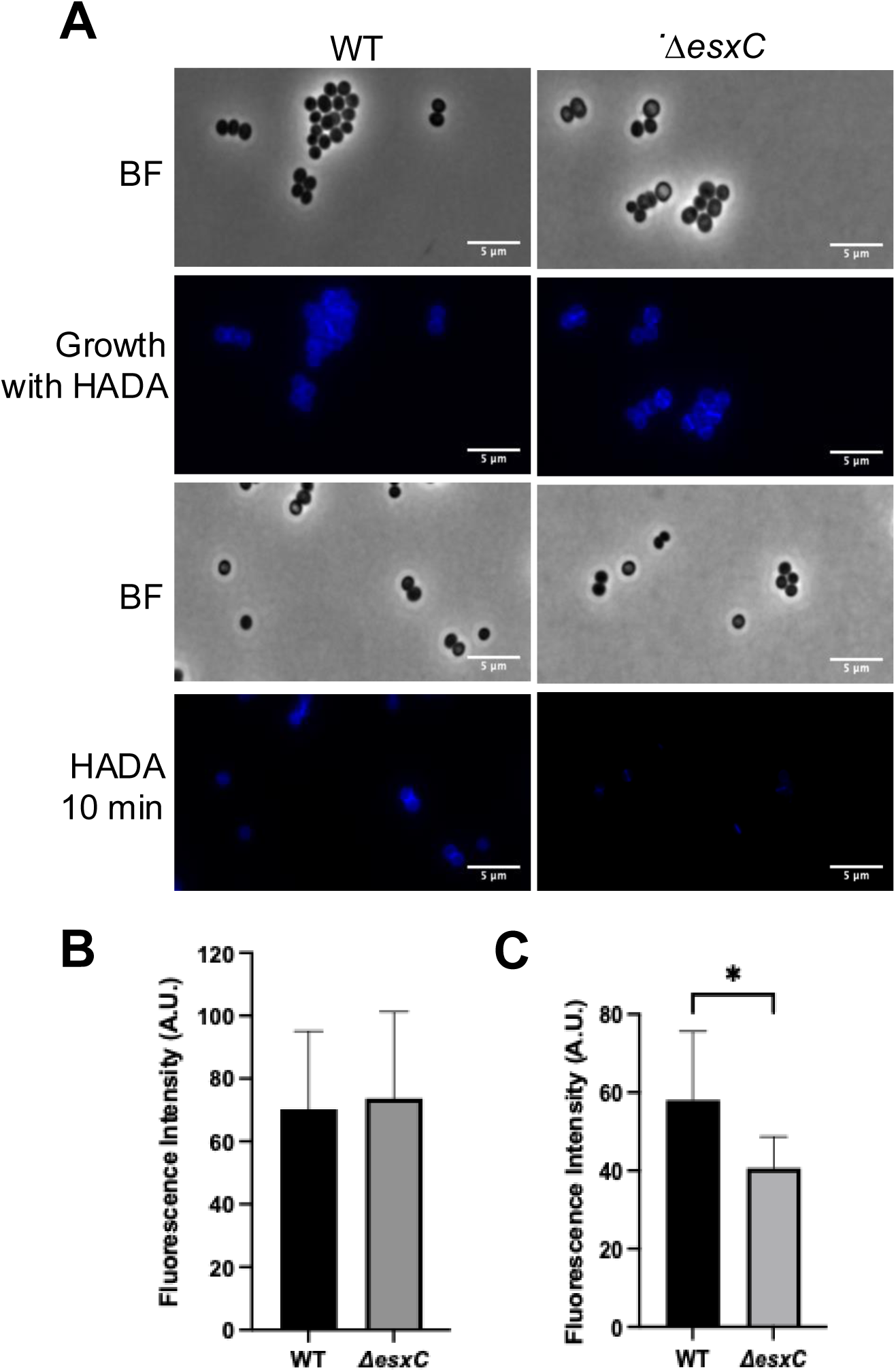
EsxC impacts cell wall synthesis. **A)** Representative widefield micrographs of *S. aureus* WT USA300 JE2 and *ΔesxC* either grown to logarithmic phase in the presence of 25 μM HADA or after 10-minute treatment with HADA once OD_600_ of 1 was reached. **B)** Quantification of fluorescence in cells grown to logarithmic phase with HADA. **C)** Quantification of fluorescence in cells treated with HADA for 10 minutes. Quantification performed on five images each from 3 independent experiments. * P< 0.01, and using an unpaired t test.

### *ΔesxC* displays an increased negative cell surface charge and increased binding of daptomycin

As we observed cell surface defects and changes to peptidoglycan synthesis, we next tested if the deletion of *esxC* could alter other cell surface properties like surface charge. A cytochrome binding assay was employed to measure surface charge; the binding of cytochrome C, a highly positively charged protein, is directly proportional to the net negative surface charge of *S. aureus* cells (27). A significantly higher level of binding of cytochrome C to the cell surface of Δ*esxC* was observed compared to the USA300 JE2 WT strain (Fig. 7 A). Similarly, Δ*esxA* also showed increased levels of binding of cytochrome C (Fig. S5). This suggests that the cell surfaces of Δ*esxC* and Δ*esxA* are more negatively charged. Much lower levels of cytochrome C bound to USA300 LAC WT and Δ*esxB* suggesting these cell surfaces are less negative (Fig. S5). CFU counts from samples after cytochrome C addition showed no differences in cell numbers and therefore it is unlikely that bacterial cell numbers influenced cytochrome C binding (Fig. 7 B, Fig S5). These data indicate that the T7SS effectors EsxC impacts the surface charge of *S. aureus*.

**Figure 7.**
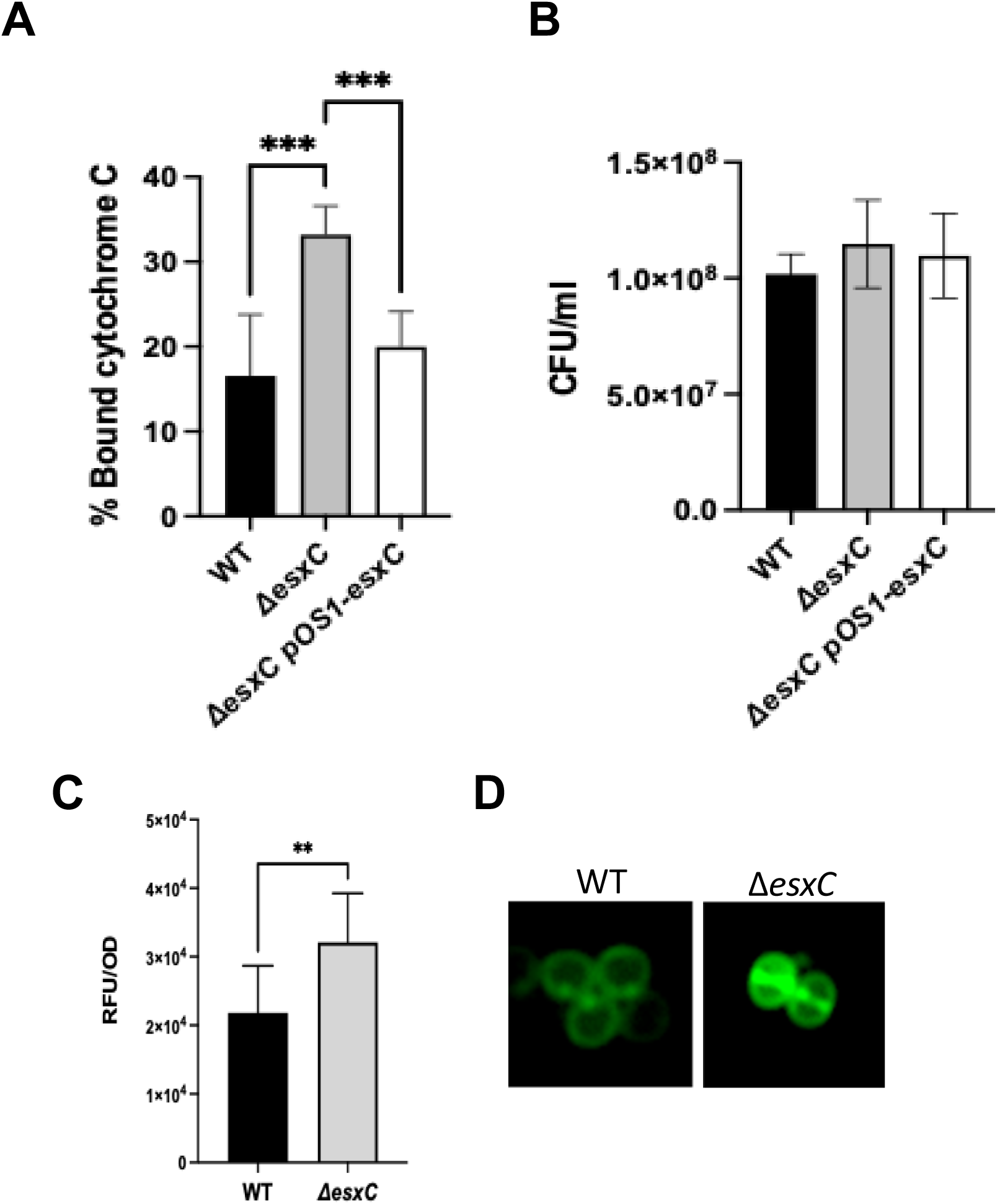
EsxC mutant binds more to daptomycin. **A-B)** Quantitative binding assay of cytochrome C to USA300 JE2 WT, Δ*esxC* and Δ*esxC* pOS1-*esxC* (A). The CFU of the starting inoculum was enumerated for each mutant (B). Mean ± SD shown, N = 3 (biological replicates), ***P < 0.001 using a one-way ANOVA. **C)** Fluorescence measurements of daptomycin-BODIPY upon binding to *S. aureus* WT USA300 JE2 and Δ*esxC.* The excitation wavelength was 488 nm and emission wavelength was 530 nm. N = 3 (biological replicates), mean + SD. P = 0.0067, using an unpaired t-test. **D)** Representative micrographs of daptomycin-BODIPY binding to *S. aureus* WT and Δ*esxC* are shown.

*S. aureus* has been reported to resist daptomycin through possessing a more positive surface charge, causing electrostatic repulsion of the positively charged daptomycin calcium complex, and thus preventing binding (28, 29). Based on the increased negative surface charge observed for the T7SS mutants in the cytochrome C assay, we hypothesised that daptomycin could bind more easily in the absence of a functional T7SS. To test this, daptomycin was tagged with BODIPY and used to quantify binding via fluorometry (Fig. 7C) and to visualise bound daptomycin by fluorescence microscopy (Fig 7D). In the presence of daptomycin, a higher level of daptomycin was observed to bind around the septum for the WT as reported previously (30). *ΔesxC* showed significantly higher fluorescence in comparison to the WT. Thus, our data suggest that reduced electrostatic repulsion of daptomycin may contribute to the increased daptomycin binding in *ΔesxC*.

### Calcium affects the growth of *ΔesxC* and sensitivity to daptomycin

Calcium is an important component in membranes, and affects both structure and charge of bacterial membranes (31). Calcium is also required for the activity of daptomycin and can enhance killing by gramicidin (32, 33). We hypothesied that EsxC may control membrane integrity through modulating the membrane binding of calcium. To test this, EsxC mutants were cultured in minimal medium with or without calcium chloride (50 mM). While both WT and mutant strains showed slow growth in minimal medium in the absence of calcium, in the presence of 50 mM calcium chloride the mutant grew much slower than the WT and the *esxC*-complemented strains (Fig 8 A,B,C). We then examined if the daptomycin toxicity was dependent on the concentration of calcium available. WT, *ΔesxC*, and the complemented strains were cultured in the presence of 5 μg/ml daptomycin and increasing calcium chloride concentrations (0 to 20 mM). In the absence of calcium, the daptomycin did not have any effect as expected. A dose-dependent decrease in growth of the WT strain was observed with increasing concentrations of calcium chloride with the WT showing slower growth at 10 mM and 20 mM CaCl_2_ compared to 1 mM (Figure 8 D, Fig S6). The mutant was more sensitive to daptomycin at all calcium concentrations compared to the WT and complemented strains, with increased growth inhibition at higher CaCl_2_ concentrations. Hence calcium binding to membranes and/or downstream signalling may be impacted in the *ΔesxC*, affecting its ability to grow in presence of calcium and its sensitivity to membrane targeting drugs like daptomycin.

**Figure 8.**
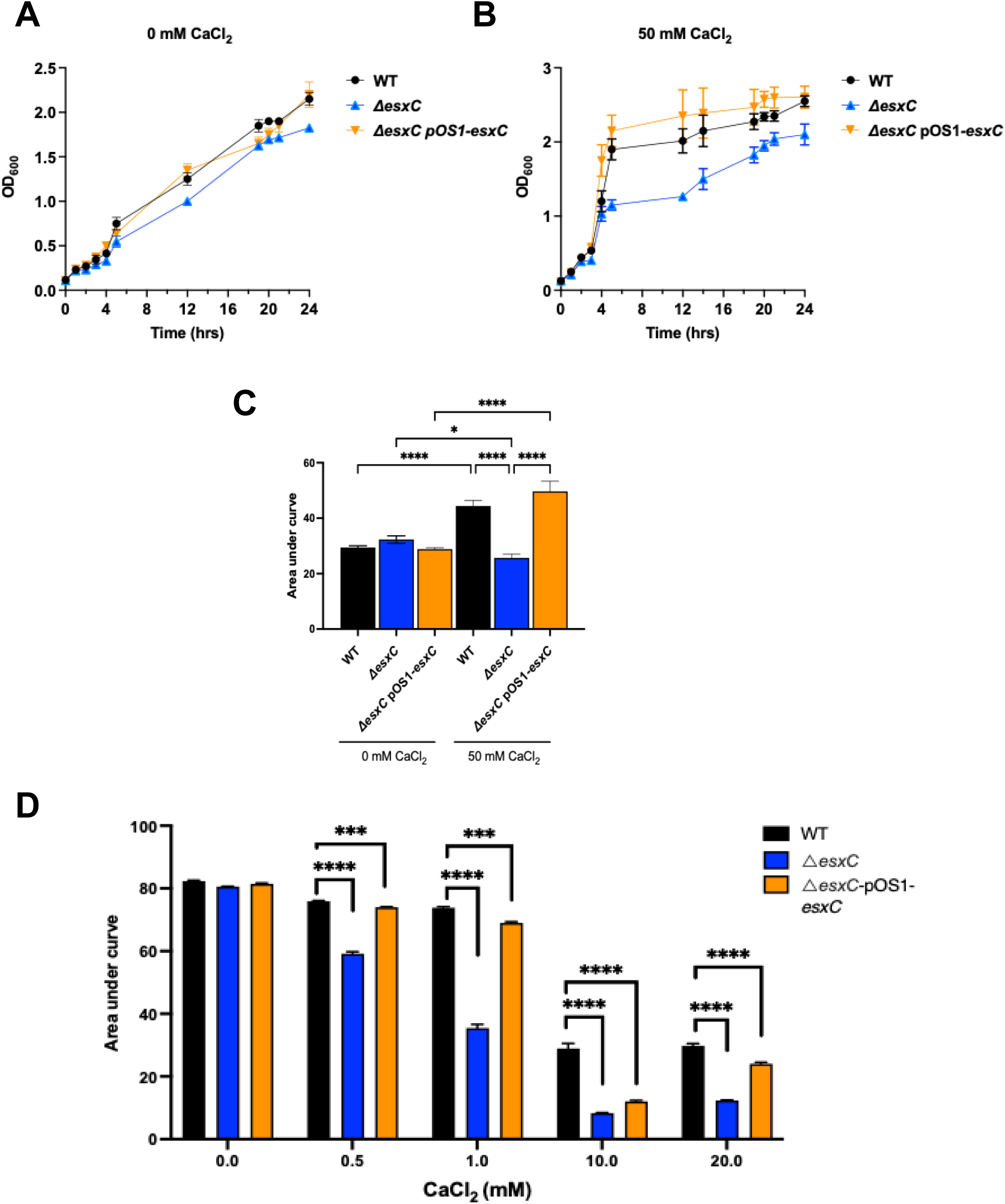
Calcium affects growth of *esxC* mutant and daptomycin sensitivity. Growth curves of *S. aureus* WT USA300 JE2, Δ*esxC*, and Δ*esxC* pOS1-*esxC* in synthetic minimal medium in the absence (A) or presence (B) of 50 mM CaCl_2_. Mean ± standard error of the mean (SEM) is shown **C)** Area under curves (AUC) were calculated from (A) and (B) and the mean AUC ± SD were plotted. **D)** *S. aureus* WT USA300 JE2, Δ*esxC*, and Δ*esxC* pOS1-*esxC* were grown in TSB supplemented with increasing calcium chloride concentrations (0 to 20 mM) in the presence of daptomycin (5 µg/ml). AUCs calculated from growth curves shown in Figure S6 are shown. Mean + SD, ***P= 0.0001, ****P <0.0001, using two-way ANOVA.

### Decreased survival of EsxC mutants during intracellular infection with daptomycin

*S. aureus* is a facultative intracellular pathogen, and daptomycin is considered to kill intracellular bacteria effectively. Hence, to determine if deletion of *esxC* from *S. aureus* would enhance intracellular killing activity, A549 human lung epithelial cells (15) were infected with *S. aureus* and treated with a high concentration of daptomycin (20 μg/ml). While there were no differences in the bacterial numbers between the WT and T7SS mutant in internalised bacteria (1 h after infection), a significant decrease was observed in the numbers of Δ*esxC* surviving within cells treated with daptomycin in comparison to WT after 4 h and 24 h after infection (Fig. 9 A). Notably, there was a significant number of WT *S. aureus* bacteria surviving intracellularly after treatment with daptomycin. Confocal microscopy imaging of infected cells also showed similar number of bacterial cells invading or internalized by mammalian cells but more *S. aureus* WT remaining 24 h after infection compared to *ΔesxC* in presence of daptomycin (Fig. 9 B, C). The differences seen in the infection assay were not due to the ability of the mutants to grow differently in DMEM-10, as strains had similar growth rates (Fig. S7A), with a growth defect in presence of daptomycin as expected (Fig. S7B). Therefore, our data suggest that daptomycin is more effective in killing intracellular *S. aureus* in the absence of T7SS.

**Figure 9.**
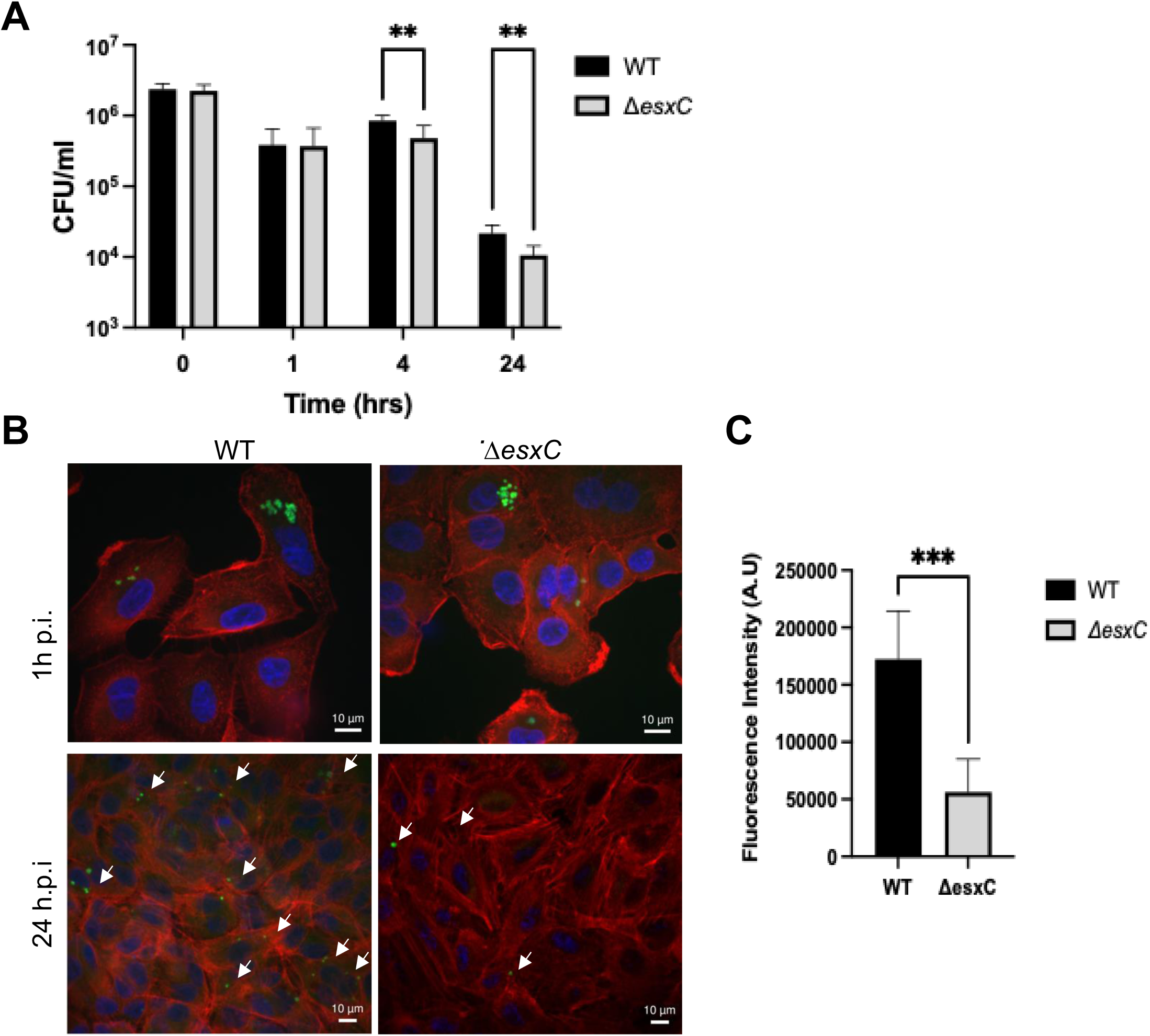
Presence of EsxC affects intracellular bacterial killing by daptomycin. **A)** CFU counts at various timepoints during epithelial cell infection assay with WT USA300 JE2, and Δ*esxC*. 0 h represents the initial inoculum, 1 h represents the CFU counts that have infected the A549 cells and 4 h and 24 h represent the survival of *S. aureus* in the presence of 20 μg/ml daptomycin, N = 3 (biological replicates), **P<0.05, two-way ANOVA. **B)** Representative confocal micrographs of cells after 1 h infection or 24 h post-infection. DAPI (blue), F-actin (red), and *S. aureus* (green, white arrows). **C)** Confocal image quantification from five images each from 3 independent experiments after 24 h. Mean ± SD of intensity, is shown, ***P<0.001 using an unpaired t-test.

**Figure 10.**
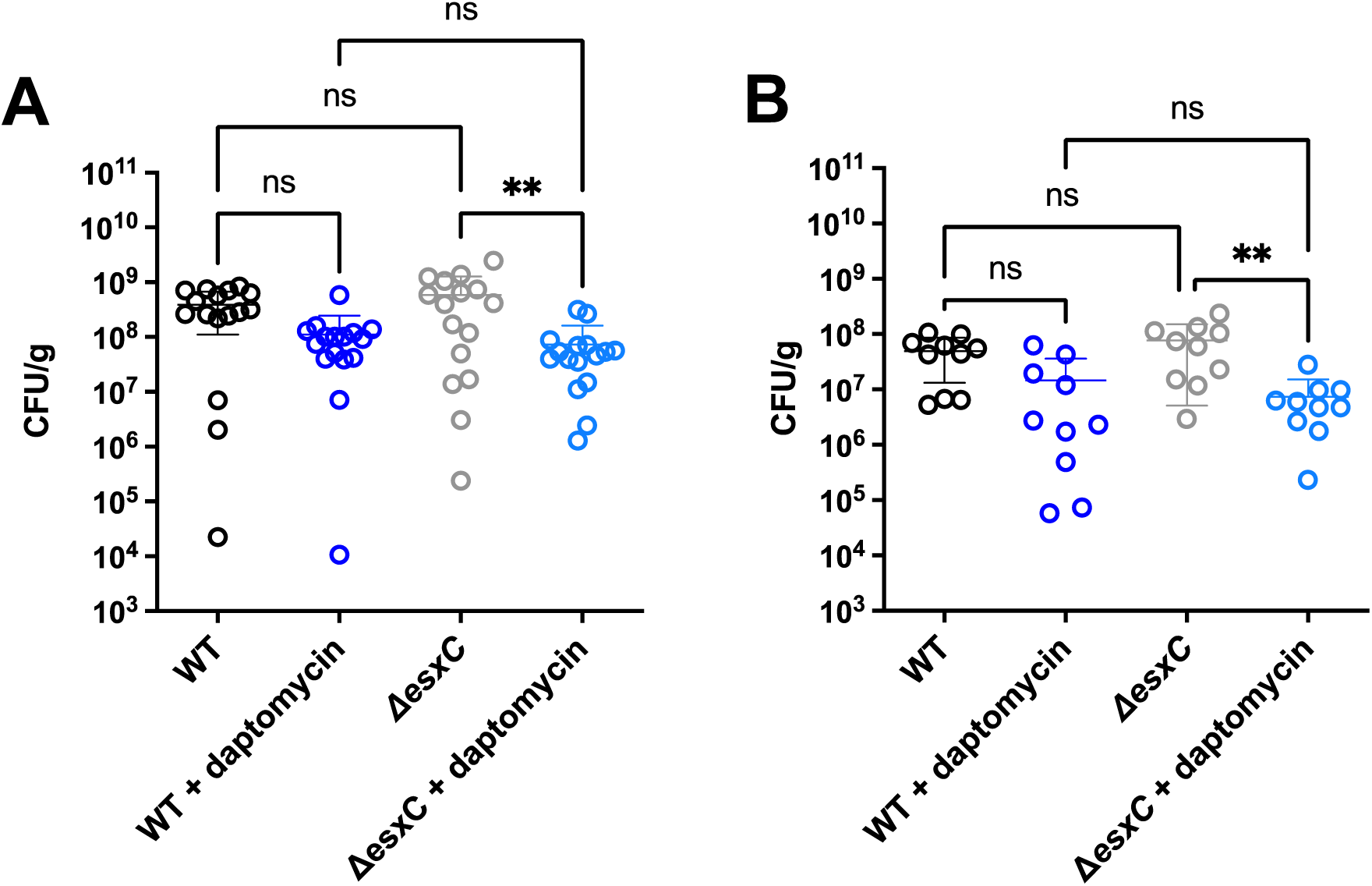
Effect of daptomycin on *esxC* mutants *in vivo*. Mice were infected in the skin with WT USA300 and Δ*esxC* mutant followed by treatment with 10 mg/kg daptomycin. **A)** CFU/g skin tissue was counted 24 h post infection (n = 8, 2 lesions/animal). Graphs show mean ± SD, P = 0.0007 using one-way ANOVA. **B)** CFU/g was calculated 72 h post infection (n = 5, 2 lesions/animal), mean ± SD shown, P = 0.0020 using a one-way ANOVA.

### *S. aureus* lacking EsxC is sensitive to daptomycin in a skin infection model

As T7SS proteins have been reported to be important in various murine infection models (6, 8, 34), we next investigated effects of daptomycin on *S. aureus* infection and the impact of EsxC on daptomycin killing *in vivo* employing a localised murine skin abscess model. 7-8-week-old BALB/c mice were infected with *S. aureus* WT and *ΔesxC* intradermally, followed by treatment (intravenous) 2 h post infection (h.p.i) and every 24 h thereafter with 10 mg/kg daptomycin. Bacterial counts were obtained from skin tissue 24 or 72 h.p.i. Weight and skin lesion diameters were recorded every day. In the non-treated groups, bacterial counts from the site of infection (lesions) showed no significant difference between WT and *ΔesxC* infected mice after 24 or 72 h of infection. A significant decrease was seen in the numbers of *ΔesxC* present in the skin of daptomycin-treated mice compared to the untreated group 24 and 72 h.p.i, while this was not the case for the WT-infected group. (Fig. 8A and B). No significant differences were observed between the areas of lesions measured for WT or *ΔesxC* after 24 h or 72 h infections and in daptomycin-treated mice (Fig S8). Although the dose of daptomycin used in the time-frame of treatment was not sufficient to treat WT *S. aureus* infection, this data suggests that the strain lacking EsxC was more susceptible to daptomycin *in vivo* compared to the WT.

## Discussion

The *S. aureus* T7SS has been strongly associated with staphylococcal virulence and the ability of this pathogen to persist during infection (7). Its importance for bacterial competition between *S. aureus* strains has been recently highlighted (13). However, the role of this system in staphylococcal physiology has been poorly defined. Here we report that EsxC, a T7SS effector is critical in maintaining cell membrane homeostasis through modulating membrane properties including fluidity, accompanied by changes in membrane protein localisation, changes to the rate of cell wall synthesis and surface charge. Our data indicate that EsxC may mediate these membrane effects through modulating calcium binding to the membrane. Consequently, EsxC impacts activity of membrane acting drugs including daptomycin, a key drug used to treat resilient staphylococcal infections. Our data demonstrate that EsxC can modulate the activity of daptomycin both intracellularly as well as during infection *in vivo*.

Several studies in mycobacterial species have linked the T7SS to cell surface integrity in different ways (19, 35–38). The ESX-1 system has been previously reported to impact cell membrane permeability in *M. tuberculosis*, although this effect appeared to be strain specific (35). However, ESX-5 which is considered essential for *in vitro* growth, has been linked to mycobacterial capsular, cell wall and outer membrane permeability (19, 36, 39). As ESX-5 was no longer essential in *M. marinum* when the membrane was permeabilised, it was proposed that ESX-5 mediates transport of cell envelope proteins to the outer membrane required for uptake of essential nutrients. While these substrates have not been identified, a substrate PPE-10 was associated with capsular integrity (19, 36). In *S. aureus,* we previously reported that the effectors, EsxC, and the ATPase, EssC, were associated with maintenance of membrane integrity in response to antimicrobial host fatty acids (20). T7SS proteins have been previously linked to membrane fluidity. Higher levels of T7SS transcription were associated with a decrease in membrane fluidity, caused by the fatty acid kinase complex, which incorporates host fatty acids into the membrane, inducing a change in membrane dynamics (24).

To explain the membrane defects associated with EsxC we considered multiple possibilities; 1. EsxC affects other membrane-associated proteins, impacting the lipid packing and fluidity of the membrane; EsxC may be a chaperone for membrane proteins and play a role in targeting them to the membrane. A recent bacterial two-hybrid library screening we performed was not successful in identifying any direct bacterial interactors, however it is possible that such interactions are transient. 2. EsxC may impact the state of the membrane by binding other molecules like metal ions required for maintaining stability of the membrane (40). 3. T7SS substrates are secreted in a co-dependent manner, so EsxC-mediated effects may be attributed to another T7SS substrate like EsxA. Our data indicate that the T7SS substrates may be involved in modulating metal binding to the membrane.

Calcium and other metal ions like magnesium are known to impact structure and function of bacterial membranes (31, 41, 42). Calcium maintains bacterial outer membrane structure through binding to bacterial lipopolysaccharide, and also influences bacterial membrane fluidity (43, 44). Calcium can also have antibacterial effects; higher concentrations of calcium can disrupt membrane integrity (42, 45, 46). Calcium affects the growth of *S. aureus* lacking EsxC indicating that the protein may modulate calcium binding to the membrane. EsxC may directly or with other T7SS effectors or membrane proteins, bind to calcium, and/ or be involved in transporting or modulating calcium binding to the cell envelope. The molecular mechanisms underlying calcium interactions of EsxC/ other T7SS substrates need further investigation.

It is also interesting that not all T7SS substrates have a cell surface associated defect which indicates substrate specificity. EsxD and EsxB did not appear to be associated with surface morphology defects by SEM as observed for EsxC and EsxA. Surface charges corelated well with daptomycin sensitivity indicating that these substrates did not play a role in the membrane homeostasis. Surface charge measurements of WT and mutants using a cytochrome C binding assay revealed that Δ*esxC* and Δ*esxA* had an increased negative surface charge, whereas *ΔesxB* did not display any differences in charge to the WT. The *esxC* and *essC* RN6390 strains however did not show similar differences by SEM, which may be due to differences in global regulatory circuits that may control cell wall synthesis (Agr, Sar) (47–49).

While *esxA* is conserved across all *S. aureus* strains, *esxC* is only found in the *essC1* locus (8, 9). It is however likely that all *S. aureus* strains encode a protein orthologous to EsxC as proteins in module 2, with variants essC2-4, have not yet been characterised and could function like EsxC. Additionally, common phenotypes were seen for strains which lacked EsxC or EsxA. Previous reports have suggested that EsxC and EsxA interact (11). Also, our previous study showed no differences in esxA levels in Δ*esxC* total cell lysates, but we find decreased EsxA in the Δ*esxC* secretome indicating co-dependent secretion of EsxA and EsxC (20). Therefore, interactions between EsxC and EsxA may be important in modulating membrane integrity.

Daptomycin is thought to bind to the cell membrane before forming a complex with cell wall biosynthesis machinery, preventing cell wall synthesis, before dispersing throughout the bacterial membrane causing depolarisation (50). Daptomycin, although not a cationic antimicrobial peptide, forms a highly positively charged complex with calcium to become active (50). Significantly more daptomycin was bound to *ΔesxC* than the WT, which supports the hypothesis that the decrease in surface charge measured in *ΔesxC* allows easier binding of daptomycin to the cell membrane. In agreement with this, T7SS substrates may influence the free calcium available for activating daptomycin. Daptomycin has also been reported to affect bacterial membrane fluidity through perturbing membrane microdomains (51). Clinical daptomycin resistant isolates have generally been associated with increased fluidity which is not associated with changes in fatty acid content (52, 53). As we show that EsxC can impact membrane fluidity, it could further contribute to daptomycin resistance by altering membrane fluidity and charge. Daptomycin resistance in *S. aureus* has been mainly associated with mutations in genes encoding for multipeptide resistance factor (MprF), a lipid biosynthetic enzyme, and the vancomycin-resistance associated sensor/regulator (VraSR), two component systems (54–56). Genome sequencing did not reveal any changes in the *ΔesxC* genome, and a proteomic analysis of mutant whole lysates did not show any modulation of proteins implicated in daptomycin resistance (20).

In contrast to daptomycin, bithionol and gramicidin are uncharged (22, 57), therefore surface charge and thus decreased electrostatic repulsion of the antibiotics is not the only factor involved in increased sensitivity of the EsxC mutant, although calcium is known to increase gramicidin activity (32). As all three antimicrobials have different mechanisms of action once bound to the membrane, before causing membrane depolarization, the subsequent steps after membrane binding appear to be important for efficacy. Therefore, daptomycin treatment which impacts both the cell membrane, and the cell wall causes a larger effect in the the EsxC mutant.

While daptomycin kills the *esxC* mutant *in vivo*, at the dose and with the strains of *S. aureus* used, there was no impact of the EsxC in bacterial survival during acute skin infection. T7SS proteins have been previously reported to be important for bacterial virulence using invasive kidney abscess, nasal colonisation and skin infection models, although studies have used different *S. aureus* strains (6, 8, 34). The lack of decrease in the lesion size of the T7SS mutant-infected mice treated with daptomycin, unlike the trend towards decreased lesion size observed for the WT (Fig S7), could indicate that the increased killing of bacteria may cause an altered local immune response. A recent study reported that in a *S. aureus* strain WU1, the T7SS was important only for invasive infection and not localized nasal colonization (58).

Currently, levels of resistance to daptomycin are low, however, with increasing resistance in other last line antibiotics, such as vancomycin, the use of daptomycin going forward will increase (59). The increased use of daptomycin will likely lead to the occurrence of more resistant isolates, therefore research into new antibiotics, and targeting new proteins that can enhance existing treatments could be a powerful approach. Our data indicate that inhibitors of T7SS proteins or of T7SS secretion could have excellent potential in combination drug therapies.

## Materials and Methods

### Bacterial strains

*S. aureus* strains used in this study are included in Table 2. Strains were cultured aerobically in tryptic soy broth (TSB) (Sigma-Aldrich) at 37°C with aeration at 180 rpm overnight. Overnight cultures were subcultured into fresh TSB and grown to a density of optical density (OD_600_) of 1, unless stated otherwise. For complemented strains TSB was supplemented with 10 μg/ml chloramphenicol.

**Table 2.** Strains and Plasmids used in this study.

| Strain or Plasmid | Description | Source or Reference |
| --- | --- | --- |
| ***Staphylococcus aureus*** |  |  |
| USA300 LAC JE2 | Plasmid-cured USA300 LAC | BEI Resources (NARSA) |
| USA300 JE2 ΔessC | USA300 LAC JE2 defective for EssC | (Kengmo Tchoupa *et al.*, 2020) |
| USA300 JE2 ΔesxC | USA300 LAC JE2 defective for EsxC | (Kengmo Tchoupa *et al.*, 2020) |
| USA300 LAC | Community acquired MRSA | (Bae and Schneewind, 2006) |
| USA300 LAC ΔesxA | USA300 LAC defective for EsxA | (Korea *et al.*, 2014) |
| USA300 LAC ΔesxB | USA300 LAC defective for EsxB | (Korea *et al.*, 2014) |
| Newman | Methicillin-sensitive *S. aureus* | (Bae and Schneewind, 2006) |
| Newman ΔesxA | Newman defective for EsxA | (Korea *et al.*, 2014) |
| Newman ΔesxB | Newman defective for EsxB | (Korea *et al.*, 2014) |
| RN6390 | NCTC8325 derivative, ΔrbsU, ΔtcaR, cured of ϕ11, ϕ12, and ϕ13 | Tracy Palmer (Kneuper *et al.*, 2014) |
| RN6390 ΔessC | RN6390 defective for EssC | Tracy Palmer (Kneuper *et al.*, 2014) |
| RN6390 ΔesxC | RN6390 defective for EsxC | Tracy Palmer (Kneuper *et al.*, 2014) |
| **Plasmids** |  |  |
| pOS1 | Empty vector for complementation | Olaf Schneewind |
| pOS1-esxC | esxC complementation vector | (Kengmo Tchoupa *et al.*, 2020) |

### Growth curves

Overnight bacterial cultures were diluted to an OD_600_ of 0.1 in TSB or TSB supplemented with antimicrobials where specified. Bacteria were grown in a 96-well plate with shaking and the OD_600_ was measured every 15 min with a FLUOstar OMEGA plate reader (BMG Labtech, UK). At appropriate time points 100 μl of the samples were taken for CFU determination. For growth curves in defined medium, overnight cultures were subcultured into minimal defined media, prepared as described by Machado and colleagues (60), at an initial OD_600_ of 0.15, and supplemented with varying concentrations of CaCl_2_ where stated. Bacterial growth was monitored manually by measuring OD_600_ at different time points using a spectrophotometer (Biochrom), due to issue with increased bacterial clumping in this medium on microtitre plates. 100 μl of the samples were also taken for CFU determination at different time points.

### Daptomycin killing assays

Bacterial cultures were grown overnight in TSB, diluted to an OD_600_ of 0.15 and grown to log phase. Bacterial cultures were then treated with 10 μg/ml of daptomycin (Acros Organics) in the presence of 1 mM CaCl_2_ (Fisher Scientific) and incubated at 37°C for a further 2 h. Aliquots were removed at indicated timepoints, serially diluted in PBS and plated on tryptic soy agar (TSA) for CFU determination.

### Scanning Electron Microscopy

Overnight bacterial cultures were diluted to an OD_600_ of 0.15 and grown to an OD_600_ of 1 in TSB. The pellet was washed with PBS twice before dropping onto poly-lysine coated 12 mm circular coverslips. Samples were fixed with 2.5% glutaraldehyde in PBS for 1 h at 4°C. The bacterial cells were dehydrated by washing with a series of ethanol solutions (20%, 50%, 70%, 90% and 100%) and dried with hexamethyldisilazane. The samples were sputter coated with carbon and then observed using a Gemini SEM 500 (Zeiss).

### Laurdan staining

Membrane fluidity was measured using the fluorescent dye Laurdan (Sigma) and was based on protocols described by Wenzel and colleagues (61). Logarithmic phase *S. aureus* was grown, and 1 ml aliquots were washed in PBS. The samples were incubated with a final concentration of 100 μM Laurdan for 5 minutes at 37°C in the dark. Samples were washed four times in pre-warmed PBS and 200 μl was transferred to a pre-warmed black walled 96 well plate in triplicate. Membrane fluidity was determined by using an excitation wavelength of 350 nm and measuring fluorescence at 460 nm and 500 nm emission wavelengths on a BioTek Cytation 5 Cell Imaging Multimode Reader (Agilent). The generalised polarisation (GP) value was calculated using the following calculation, GP = (I_460_-I_500_) / (I_460_+I_500_).

### Pyrene decanoic acid staining

O/N bacterial cultures were diluted to an OD_600_ of 0.15 in TSB and were grown to an OD_600_ of 1.0. Bacteria were washed with PBS prior to treatment for 30 min at 37°C with 37.5 µg/mL lysostaphin in PBS containing 20% sucrose. The spheroblasts were then centrifuged at 8000 × g for 10 min, and the pellet resuspended in the labelling solution (PBS, 20% sucrose, 0.01% F-127, 5 µM pyrene decanoic acid). The incubation in the dark was done for 1h at 25°C under gentle rotation. PBS supplemented with 20% sucrose was used to wash the stained spheroblasts that were afterwards transferred to 96-well plates for fluorescence 450 nm measurements as previously described (24).

### DiIC12 staining

O/N bacterial cultures were diluted to an OD_600_ of 0.15 in TSB with 1 μg/mL 1,1′-didodecyl-3,3,3′,3′-tetramethylindocarbocyanine perchlorate (DiIC12) (Invitrogen). Cultures were grown to an OD_600_ of 1.0, centrifuged and washed twice with fresh TSB. Samples were spotted on to agarose pads and imaged using a Leica DMi8 widefield microscope (Leica Microsystems, UK). Acquired images were analysed with the ImageJ processing package, Fiji.

### Cytochrome C binding assay

Overnight bacterial cultures were pelleted and washed with 20 mM MOPS buffer, pH 7. The cells were resuspended in MOPS buffer at an OD_600_ of 1 and incubated with 50 μg/ml cytochrome C at room temperature for 15 min. The cells were pelleted and the remaining cytochrome C in the supernatant was measured spectrophotometrically at OD_410_ using a FLUOstar OMEGA plate reader (BMG Labtech, UK).

### HADA incorporation assay

Peptidoglycan synthesis was investigated by measuring HCC-amino-D-alanine (HADA) incorporation. Overnight *S. aureus* cultures were diluted into TSB supplemented with 25 μM HADA and incubated at 37°C in the dark. When OD_600_ of 1 was reached cultures were washed three times and fixed with 4% PFA. Alternatively, cultures were treated with daptomycin for 30 min. before washing and fixing. Microscopy was undertaken on these samples using the DAPI filter set on a Leica DMi8 widefield microscope.

### Daptomycin BODIPY binding assay

*S. aureus* was grown as previously described and treated with 1μg/ml daptomycin-BODIPY once logarithmic phase had been reached. The samples were incubated at 37°C for 30 min in the dark, then washed with PBS three times. 200 μl were aliquoted into black 96 well plates in triplicate and fluorescence was measured using a BioTek Cytation 5 Cell Imaging Multimode Reader (Agilent). The excitation wavelength used was 488 nm and emission wavelength used was 530 nm.

### Invasion and survival assays

Infection assays were carried out in 24 well tissue culture treated plates (Falcon), or RS treated glass chamber slides (154526, Nunc, Lab Tek II). Overnight *S. aureus* cultures were diluted to an OD_600_ of 0.15 and grown to an OD_600_ of 1. Cultures were resuspended in DMEM media to a density of 2.5 x 10^6^ CFU/ml. A549 epithelial cells were seeded in 24 well plates at a density of 2.5 x 10^5^ one day before being infected with *S. aureus* at a multiplicity of infection (MOI) of 10. Cells were infected for 1 h at 37°C with 5 % CO_2_. To kill any extracellular bacteria, cells were washed with DMEM media containing 20 μg/ml lysostaphin (Sigma-Aldrich) and 50 μg/ml gentamycin (Melford) and incubated for a further 30 minutes. Cells were washed twice with PBS, lysed with 1 ml cold sterile water and vigorous pipetting before plating for CFU determination. To investigate the course of infection after extracellular bacterial killing, the cells are washed twice with PBS and refreshed with DMEM with 1 mM CaCl_2_ only, or DMEM supplemented with 1 mM CaCl_2_ and 10 μg/ml daptomycin and incubated until the desired timepoint. The above protocol to lyse the cells is then followed.

### Murine skin abscess model

All animal procedures were carried out as per protocols in the Project Licence PCEC27E7D. All local ethical and home office approvals were obtained for animal experimentation. For the murine skin abscess model, female 7/8-week-old BALB/c mice (Charles River Laboratories, UK) flanks were shaved before injection, and inoculated through an intradermal route with 50 μl of bacterial suspension per flank (one injection on each flank), resulting in a final bacterial count of 1-2 x 10^6^ per mouse. Overnight *S. aureus* cultures were subcultured, grown to OD_600_ of 1, centrifuged and washed twice with PBS, and diluted in PBS to achieve a final CFU/ml of 1-2 x 10^7^ for infection. 2 h after infection mice were given an intravenous dose of daptomycin or PBS via tail vein injection. This was repeated once every 24 h. During the experiment, mice were weighed daily, and any abscess formation was measured and recorded. After 24 h or 72 h mice were euthanised by CO_2_ inhalation and skin at the infection site was incised. The skin samples were weighed and homogenised in PBS using a FastPrep (MP Biomedicals), at 4 m/s for 60 seconds for a total of 6 cycles. Homogenates were diluted in PBS and CFU/ml was determined. CFU counts were normalized to lesion weight.

### Statistical Analysis

All experiments were performed at least three independent times. Statistical analysis was performed using GraphPad Prism 9.0 (MacOS version 9.5.1). Pairwise statistical analysis was carried out by the Mann-Whitney U test or t test? for two groups, or One way ANOVAs for multiple groups, followed by post-hoc Tukey’s HSD tests where necessary. Significance was denoted by asterisks; * = P < 0.05, ** = P < 0.01, *** = P < 0.001, **** = P < 0.0001 and ns = not significant.

## Acknowledgements

We would like to acknowledge funding from MIBTP (grant 2097375) to Victoria Smith and the Medical Research Council grants (MR/N010140/1 and MR/X00161X/1) for funding.

We would like to thank Novartis Vaccines, Siena (GSK biologicals) Italy for providing the *ΔesxA* and *ΔesxB* strains, and Prof Tracy Palmer for providing the RN6390 *ΔesxC* and *ΔessC* strains, used in this study. We acknowledge the Midlands Regional Cryo-EM Facility, hosted at the Warwick Advanced Bioimaging Research Technology Platform, for the use of the JEOL 2100Plus.

## Author contributions

VS, AKT and MU conceptualised ideas, designed experiments, VS, GC, RF, KEW, RGM, JY, AKT conducted experiments, MU and SP provided resources for the study, VS, AKT, GC, KEW, RGM, RF and MU analysed data, AKT and MU supervised study, MU, VS, RF and AKT drafted manuscript and all authors were involved in reviewing manuscript draft.

## References

1. Tong SY, Davis JS, Eichenberger E, Holland TL, Fowler VG, Jr. 2015. Staphylococcus aureus infections: epidemiology, pathophysiology, clinical manifestations, and management. Clin Microbiol Rev 28:603–61.

2. Chambers HF, Deleo FR. 2009. Waves of resistance: Staphylococcus aureus in the antibiotic era. Nat Rev Microbiol 7:629–41.

3. Lee BY, Singh A, David MZ, Bartsch SM, Slayton RB, Huang SS, Zimmer SM, Potter MA, Macal CM, Lauderdale DS, Miller LG, Daum RS. 2013. The economic burden of community-associated methicillin-resistant Staphylococcus aureus (CA-MRSA). Clin Microbiol Infect 19:528–36.

4. Cheung GYC, Bae JS, Otto M. 2021. Pathogenicity and virulence of Staphylococcus aureus. Virulence 12:547–569.

5. Bottai D, Groschel MI, Brosch R. 2016. Type VII Secretion Systems in Gram-Positive Bacteria. Curr Top Microbiol Immunol doi:10.1007/82_2015_5015.

6. Burts ML, Williams WA, DeBord K, Missiakas DM. 2005. EsxA and EsxB are secreted by an ESAT-6-like system that is required for the pathogenesis of Staphylococcus aureus infections. Proc Natl Acad Sci U S A 102:1169–74.

7. Unnikrishnan M, Constantinidou C, Palmer T, Pallen MJ. 2017. The Enigmatic Esx Proteins: Looking Beyond Mycobacteria. Trends Microbiol 25:192–204.

8. Kneuper H, Cao ZP, Twomey KB, Zoltner M, Jager F, Cargill JS, Chalmers J, van der Kooi-Pol MM, van Dijl JM, Ryan RP, Hunter WN, Palmer T. 2014. Heterogeneity in ess transcriptional organization and variable contribution of the Ess/Type VII protein secretion system to virulence across closely related Staphylocccus aureus strains. Mol Microbiol 93:928–43.

9. Warne B, Harkins CP, Harris SR, Vatsiou A, Stanley-Wall N, Parkhill J, Peacock SJ, Palmer T, Holden MT. 2016. The Ess/Type VII secretion system of Staphylococcus aureus shows unexpected genetic diversity. BMC Genomics 17:222.

10. Bowman L, Palmer T. 2021. The Type VII Secretion System of Staphylococcus. Annu Rev Microbiol 75:471–494.

11. Anderson M, Aly KA, Chen YH, Missiakas D. 2013. Secretion of atypical protein substrates by the ESAT-6 secretion system of Staphylococcus aureus. Mol Microbiol 90:734–43.

12. Mielich-Suss B, Wagner RM, Mietrach N, Hertlein T, Marincola G, Ohlsen K, Geibel S, Lopez D. 2017. Flotillin scaffold activity contributes to type VII secretion system assembly in Staphylococcus aureus. PLoS Pathog 13:e1006728.

13. Cao Z, Casabona MG, Kneuper H, Chalmers JD, Palmer T. 2016. The type VII secretion system of Staphylococcus aureus secretes a nuclease toxin that targets competitor bacteria. Nat Microbiol 2:16183.

14. Ulhuq FR, Gomes MC, Duggan GM, Guo M, Mendonca C, Buchanan G, Chalmers JD, Cao Z, Kneuper H, Murdoch S, Thomson S, Strahl H, Trost M, Mostowy S, Palmer T. 2020. A membrane-depolarizing toxin substrate of the Staphylococcus aureus type VII secretion system mediates intraspecies competition. Proc Natl Acad Sci U S A 117:20836–20847.

15. Korea CG, Balsamo G, Pezzicoli A, Merakou C, Tavarini S, Bagnoli F, Serruto D, Unnikrishnan M. 2014. Staphylococcal Esx proteins modulate apoptosis and release of intracellular Staphylococcus aureus during infection in epithelial cells. Infect Immun 82:4144–53.

16. Anderson M, Ohr RJ, Aly KA, Nocadello S, Kim HK, Schneewind CE, Schneewind O, Missiakas D. 2017. EssE Promotes Staphylococcus aureus ESS-Dependent Protein Secretion To Modify Host Immune Responses during Infection. J Bacteriol 199.

17. Fyans JK, Bignell D, Loria R, Toth I, Palmer T. 2013. The ESX/type VII secretion system modulates development, but not virulence, of the plant pathogen Streptomyces scabies. Mol Plant Pathol 14:119–30.

18. Flint JL, Kowalski JC, Karnati PK, Derbyshire KM. 2004. The RD1 virulence locus of Mycobacterium tuberculosis regulates DNA transfer in Mycobacterium smegmatis. Proc Natl Acad Sci U S A 101:12598–603.

19. Ates LS, Ummels R, Commandeur S, van de Weerd R, Sparrius M, Weerdenburg E, Alber M, Kalscheuer R, Piersma SR, Abdallah AM, Abd El Ghany M, Abdel-Haleem AM, Pain A, Jimenez CR, Bitter W, Houben EN. 2015. Essential Role of the ESX-5 Secretion System in Outer Membrane Permeability of Pathogenic Mycobacteria. PLoS Genet 11:e1005190.

20. Kengmo Tchoupa A, Watkins KE, Jones RA, Kuroki A, Alam MT, Perrier S, Chen Y, Unnikrishnan M. 2020. The type VII secretion system protects Staphylococcus aureus against antimicrobial host fatty acids. Sci Rep 10:14838.

21. Huang HW. 2020. DAPTOMYCIN, its membrane-active mechanism vs. that of other antimicrobial peptides. Biochim Biophys Acta Biomembr 1862:183395.

22. Kim W, Zou G, Hari TPA, Wilt IK, Zhu W, Galle N, Faizi HA, Hendricks GL, Tori K, Pan W, Huang X, Steele AD, Csatary EE, Dekarske MM, Rosen JL, Ribeiro NQ, Lee K, Port J, Fuchs BB, Vlahovska PM, Wuest WM, Gao H, Ausubel FM, Mylonakis E. 2019. A selective membrane-targeting repurposed antibiotic with activity against persistent methicillin-resistant Staphylococcus aureus. Proc Natl Acad Sci U S A 116:16529–16534.

23. Strahl H, Burmann F, Hamoen LW. 2014. The actin homologue MreB organizes the bacterial cell membrane. Nat Commun 5:3442.

24. Lopez MS, Tan IS, Yan D, Kang J, McCreary M, Modrusan Z, Austin CD, Xu M, Brown EJ. 2017. Host-derived fatty acids activate type VII secretion in Staphylococcus aureus. Proc Natl Acad Sci U S A 114:11223–11228.

25. Scheinpflug K, Krylova O, Strahl H. 2017. Measurement of Cell Membrane Fluidity by Laurdan GP: Fluorescence Spectroscopy and Microscopy. Methods Mol Biol 1520:159–174.

26. Kuru E, Hughes HV, Brown PJ, Hall E, Tekkam S, Cava F, de Pedro MA, Brun YV, VanNieuwenhze MS. 2012. In Situ probing of newly synthesized peptidoglycan in live bacteria with fluorescent D-amino acids. Angew Chem Int Ed Engl 51:12519–23.

27. Peschel A, Otto M, Jack RW, Kalbacher H, Jung G, Gotz F. 1999. Inactivation of the dlt operon in Staphylococcus aureus confers sensitivity to defensins, protegrins, and other antimicrobial peptides. J Biol Chem 274:8405–10.

28. Bertsche U, Yang SJ, Kuehner D, Wanner S, Mishra NN, Roth T, Nega M, Schneider A, Mayer C, Grau T, Bayer AS, Weidenmaier C. 2013. Increased cell wall teichoic acid production and D-alanylation are common phenotypes among daptomycin-resistant methicillin-resistant Staphylococcus aureus (MRSA) clinical isolates. PLoS One 8:e67398.

29. Mishra NN, Bayer AS, Weidenmaier C, Grau T, Wanner S, Stefani S, Cafiso V, Bertuccio T, Yeaman MR, Nast CC, Yang SJ. 2014. Phenotypic and genotypic characterization of daptomycin-resistant methicillin-resistant Staphylococcus aureus strains: relative roles of mprF and dlt operons. PLoS One 9:e107426.

30. Monteiro JM, Fernandes PB, Vaz F, Pereira AR, Tavares AC, Ferreira MT, Pereira PM, Veiga H, Kuru E, VanNieuwenhze MS, Brun YV, Filipe SR, Pinho MG. 2015. Cell shape dynamics during the staphylococcal cell cycle. Nat Commun 6:8055.

31. Smith RJ. 1995. Calcium and bacteria. Adv Microb Physiol 37:83–133.

32. Fang ST, Huang SH, Yang CH, Liou JW, Mani H, Chen YC. 2022. Effects of Calcium Ions on the Antimicrobial Activity of Gramicidin A. Biomolecules 12.

33. Jung D, Rozek A, Okon M, Hancock RE. 2004. Structural transitions as determinants of the action of the calcium-dependent antibiotic daptomycin. Chem Biol 11:949–57.

34. Wang Y, Hu M, Liu Q, Qin J, Dai Y, He L, Li T, Zheng B, Zhou F, Yu K, Fang J, Liu X, Otto M, Li M. 2016. Role of the ESAT-6 secretion system in virulence of the emerging community-associated Staphylococcus aureus lineage ST398. Sci Rep 6:25163.

35. Garces A, Atmakuri K, Chase MR, Woodworth JS, Krastins B, Rothchild AC, Ramsdell TL, Lopez MF, Behar SM, Sarracino DA, Fortune SM. 2010. EspA acts as a critical mediator of ESX1-dependent virulence in Mycobacterium tuberculosis by affecting bacterial cell wall integrity. PLoS Pathog 6:e1000957.

36. Ates LS, van der Woude AD, Bestebroer J, van Stempvoort G, Musters RJ, Garcia-Vallejo JJ, Picavet DI, Weerd R, Maletta M, Kuijl CP, van der Wel NN, Bitter W. 2016. The ESX-5 System of Pathogenic Mycobacteria Is Involved In Capsule Integrity and Virulence through Its Substrate PPE10. PLoS Pathog 12:e1005696.

37. Nath Y, Ray SK, Buragohain AK. 2018. Essential role of the ESX-3 associated eccD3 locus in maintaining the cell wall integrity of Mycobacterium smegmatis. Int J Med Microbiol 308:784–795.

38. Bosserman RE, Champion PA. 2017. Esx Systems and the Mycobacterial Cell Envelope: What’s the Connection? J Bacteriol 199.

39. Bottai D, Di Luca M, Majlessi L, Frigui W, Simeone R, Sayes F, Bitter W, Brennan MJ, Leclerc C, Batoni G, Campa M, Brosch R, Esin S. 2012. Disruption of the ESX-5 system of Mycobacterium tuberculosis causes loss of PPE protein secretion, reduction of cell wall integrity and strong attenuation. Mol Microbiol 83:1195–209.

40. Thomas KJ, 3rd, Rice CV. 2014. Revised model of calcium and magnesium binding to the bacterial cell wall. Biometals 27:1361–70.

41. Pedersen UR, Leidy C, Westh P, Peters GH. 2006. The effect of calcium on the properties of charged phospholipid bilayers. Biochim Biophys Acta 1758:573–82.

42. Melcrova A, Pokorna S, Pullanchery S, Kohagen M, Jurkiewicz P, Hof M, Jungwirth P, Cremer PS, Cwiklik L. 2016. The complex nature of calcium cation interactions with phospholipid bilayers. Sci Rep 6:38035.

43. Ginez LD, Osorio A, Vazquez-Ramirez R, Arenas T, Mendoza L, Camarena L, Poggio S. 2022. Changes in fluidity of the E. coli outer membrane in response to temperature, divalent cations and polymyxin-B show two different mechanisms of membrane fluidity adaptation. FEBS J 289:3550–3567.

44. Schindler M, Osborn MJ. 1979. Interaction of divalent cations and polymyxin B with lipopolysaccharide. Biochemistry 18:4425–30.

45. Xie Y, Yang L. 2016. Calcium and Magnesium Ions Are Membrane-Active against Stationary-Phase Staphylococcus aureus with High Specificity. Sci Rep 6:20628.

46. Asif A, Mohsin H, Tanvir R, Rehman Y. 2017. Revisiting the Mechanisms Involved in Calcium Chloride Induced Bacterial Transformation. Front Microbiol 8:2169.

47. Fujimoto DF, Bayles KW. 1998. Opposing roles of the Staphylococcus aureus virulence regulators, Agr and Sar, in Triton X-100- and penicillin-induced autolysis. J Bacteriol 180:3724–6.

48. Cheung GY, Wang R, Khan BA, Sturdevant DE, Otto M. 2011. Role of the accessory gene regulator agr in community-associated methicillin-resistant Staphylococcus aureus pathogenesis. Infect Immun 79:1927–35.

49. Zielinska AK, Beenken KE, Joo HS, Mrak LN, Griffin LM, Luong TT, Lee CY, Otto M, Shaw LN, Smeltzer MS. 2011. Defining the strain-dependent impact of the Staphylococcal accessory regulator (sarA) on the alpha-toxin phenotype of Staphylococcus aureus. J Bacteriol 193:2948–58.

50. Grein F, Muller A, Scherer KM, Liu X, Ludwig KC, Klockner A, Strach M, Sahl HG, Kubitscheck U, Schneider T. 2020. Ca(2+)-Daptomycin targets cell wall biosynthesis by forming a tripartite complex with undecaprenyl-coupled intermediates and membrane lipids. Nat Commun 11:1455.

51. Muller A, Wenzel M, Strahl H, Grein F, Saaki TNV, Kohl B, Siersma T, Bandow JE, Sahl HG, Schneider T, Hamoen LW. 2016. Daptomycin inhibits cell envelope synthesis by interfering with fluid membrane microdomains. Proc Natl Acad Sci U S A 113:E7077–E7086.

52. Jones T, Yeaman MR, Sakoulas G, Yang SJ, Proctor RA, Sahl HG, Schrenzel J, Xiong YQ, Bayer AS. 2008. Failures in clinical treatment of Staphylococcus aureus Infection with daptomycin are associated with alterations in surface charge, membrane phospholipid asymmetry, and drug binding. Antimicrob Agents Chemother 52:269–78.

53. Mishra NN, Bayer AS. 2013. Correlation of cell membrane lipid profiles with daptomycin resistance in methicillin-resistant Staphylococcus aureus. Antimicrob Agents Chemother 57:1082–5.

54. Ernst CM, Peschel A. 2019. MprF-mediated daptomycin resistance. Int J Med Microbiol 309:359–363.

55. Mehta S, Cuirolo AX, Plata KB, Riosa S, Silverman JA, Rubio A, Rosato RR, Rosato AE. 2012. VraSR two-component regulatory system contributes to mprF-mediated decreased susceptibility to daptomycin in in vivo-selected clinical strains of methicillin-resistant Staphylococcus aureus. Antimicrob Agents Chemother 56:92–102.

56. Nguyen AH, Hood KS, Mileykovskaya E, Miller WR, Tran TT. 2022. Bacterial cell membranes and their role in daptomycin resistance: A review. Front Mol Biosci 9:1035574.

57. Rostovtseva TK, Aguilella VM, Vodyanoy I, Bezrukov SM, Parsegian VA. 1998. Membrane surface-charge titration probed by gramicidin A channel conductance. Biophys J 75:1783–92.

58. Bobrovskyy M, Chen X, Missiakas D. 2023. The Type 7b Secretion System of S. aureus and Its Role in Colonization and Systemic Infection. Infect Immun 91:e0001523.

59. Shariati A, Dadashi M, Moghadam MT, van Belkum A, Yaslianifard S, Darban-Sarokhalil D. 2020. Global prevalence and distribution of vancomycin resistant, vancomycin intermediate and heterogeneously vancomycin intermediate Staphylococcus aureus clinical isolates: a systematic review and meta-analysis. Sci Rep 10:12689.

60. Machado H, Weng LL, Dillon N, Seif Y, Holland M, Pekar JE, Monk JM, Nizet V, Palsson BO, Feist AM. 2019. Strain-Specific Metabolic Requirements Revealed by a Defined Minimal Medium for Systems Analyses of Staphylococcus aureus. Appl Environ Microbiol 85.

61. Wenzel M, Vischer NOE, Strahl H, Hamoen LW. 2018. Assessing Membrane Fluidity and Visualizing Fluid Membrane Domains in Bacteria Using Fluorescent Membrane Dyes. Bio Protoc 8:e3063.

