## supplemental table for "The staphylococcal type VII secretion system effector EsxC impacts daptomycin sensitivity through controlling bacterial cell envelope integrity"

**Table S1. Secreted proteins significantly changed in Δ*esxC* relative to WT USA300 JE2.**

| Uniprot ID | $log_2$ fold change | Adjusted *P* value | Description |
|---|---|---|---|
| A0A0H2XIV9 | -4.27 | 7.11E-10 | T7SS protein EssD |
| A0A0H2XHT7 | -2.78 | 7.96E-09 | Phosphocarrier protein HPr |
| A0A0H2XH98 | -1.75 | 1.34E-07 | Hydrolase, MutT/nudix family |
| A0A0H2XIK2 | -1.71 | 1.34E-07 | T7SS protein EsxC |
| A0A0H2XI99 | -1.45 | 0.003045 | T7SS protein EsxA |
| A0A0H2XFA5 | -1.28 | 4.03E-05 | Putative GTP-binding protein |
| A0A0H2XEF1 | 1.55 | 3.40E-06 | 2-oxoisovalerate dehydrogenase |
| Q2FEF1 | 1.63 | 0.000836 | Isopentenyl-diphosphate delta-isomerase |
| A0A0H2XKB6 | 1.90 | 0.00012 | Uncharacterized protein |
| A0A0H2XH53 | 2.58 | 0.042723 | Uncharacterized protein |
