## supplemental figures for "The staphylococcal type VII secretion system effector EsxC impacts daptomycin sensitivity through controlling bacterial cell envelope integrity"

Fig S1

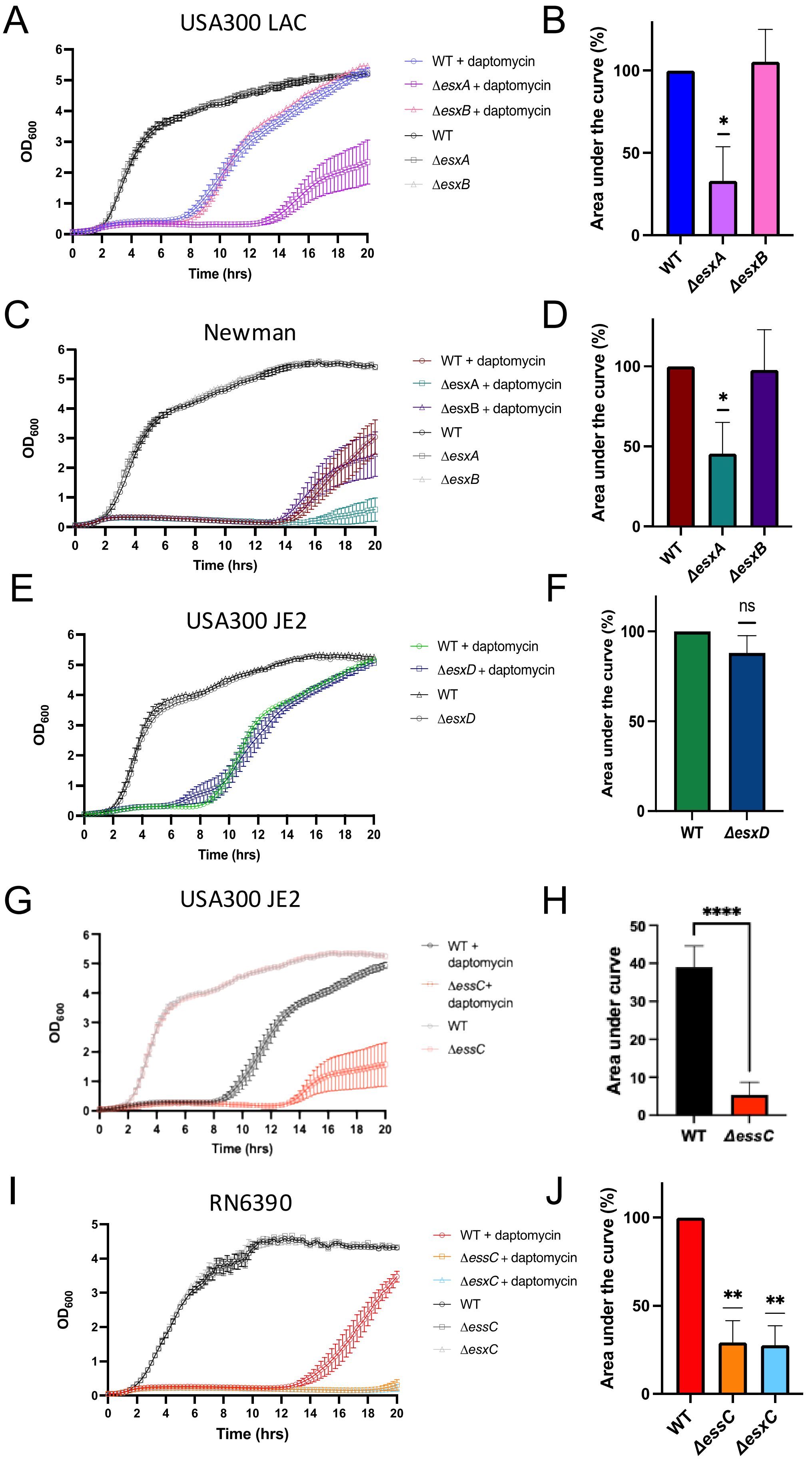


**Figure S1.** Growth curves in TSB for A) *S. aureus* USA300 LAC Δ*esxA* and Δ*esxB*, C) *S. aureus* Newman Δ*esxA* and Δ*esxB*, E) *S. aureus* USA300 JE2 WT and Δ*esxD*, G) *S. aureus* USA300 JE2 WT and Δ*essC* I) *S. aureus* RN6390 WT, Δ*essC* and Δ*esxC*, in the absence or presence of 5 μg/ml daptomycin and 1 mM $CaCl_2$. Mean ± SEM are shown, N = 3. The AUC was calculated for the growth curves of all strains cultured in presence of daptomycin in A, C, E and G and I are presented as % relative to WT, mean + SD is shown (B, D, F, H, J). * P ≤ 0.05, ** P ≤ 0.01 using a one-sample t test. J) shows AUC of I) ****P ≤ 0.0001, unpaired t test

Fig S2

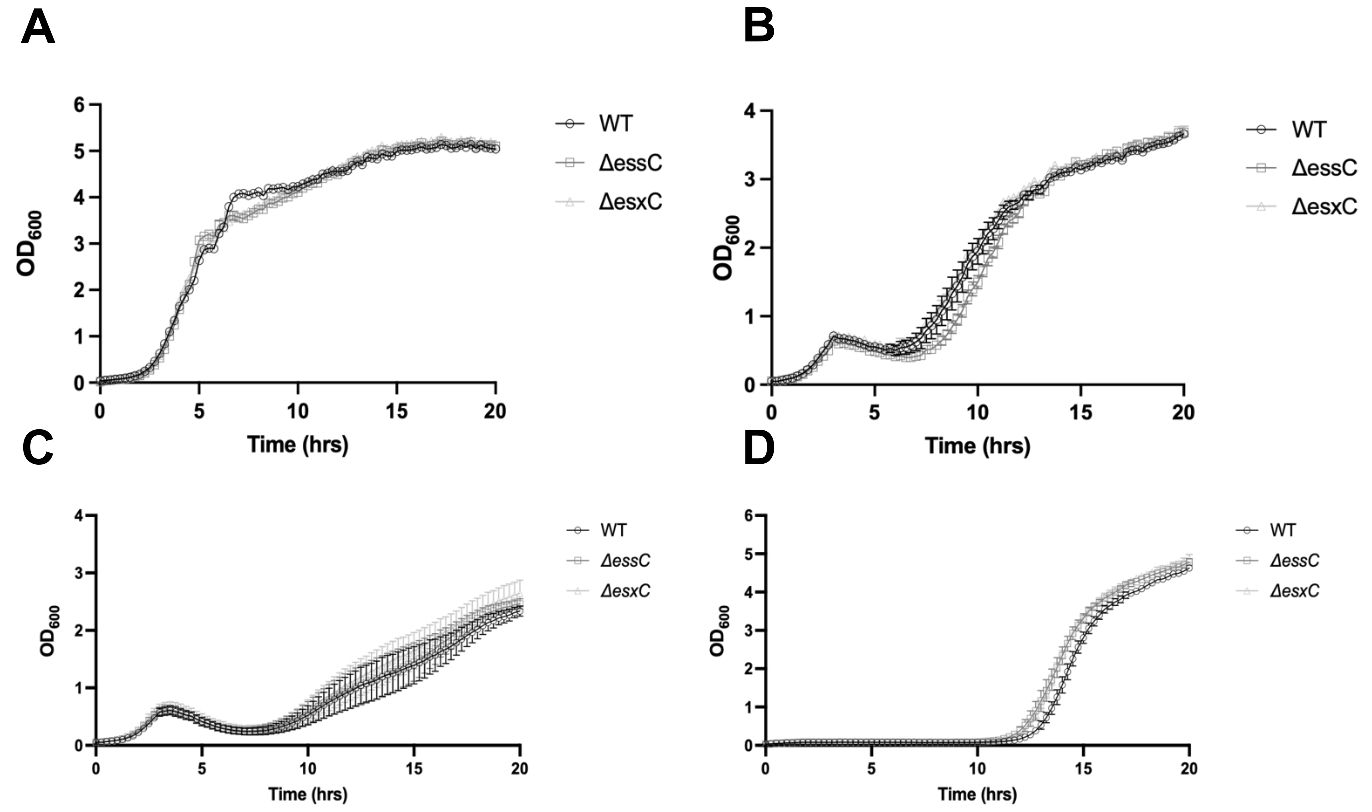


**Figure S2**. Growth curves of USA300 JE2 WT, *ΔessC* and Δ*esxC* in the presence of (A) 4 µg/ml ciprofloxacin (B) 0.5 µg/ml mitomycin C. (C) 1 µg/ml oxacillin. (D) 8 µg/ml vancomycin. For all growth curves mean and SEM are shown, N = 3 (biological replicates).

Fig S3

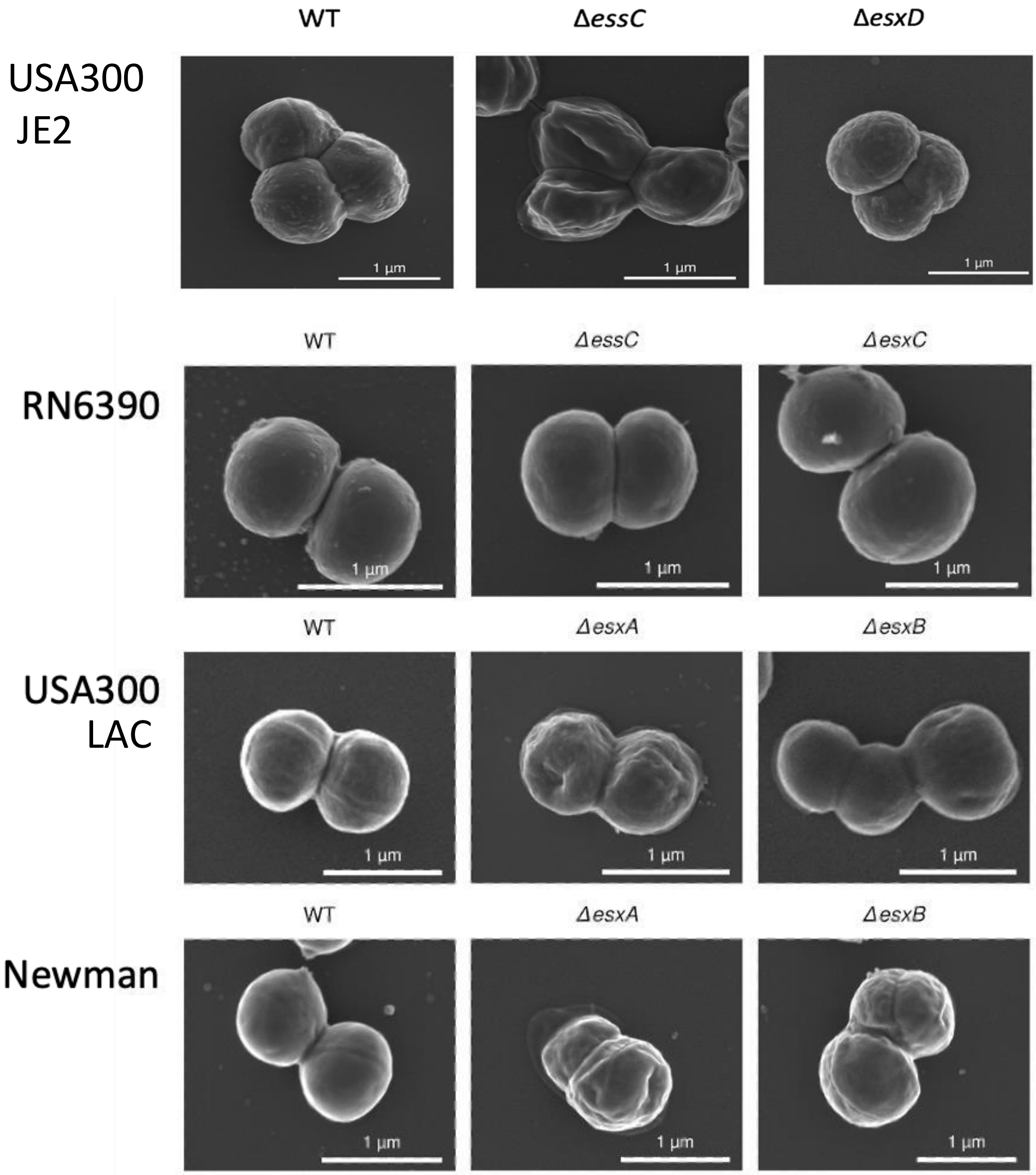


**Figure S3.** Scanning electron micrographs of *S. aureus* RN6390, USA300, Newman and their T7SS mutants**.**

Fig S4

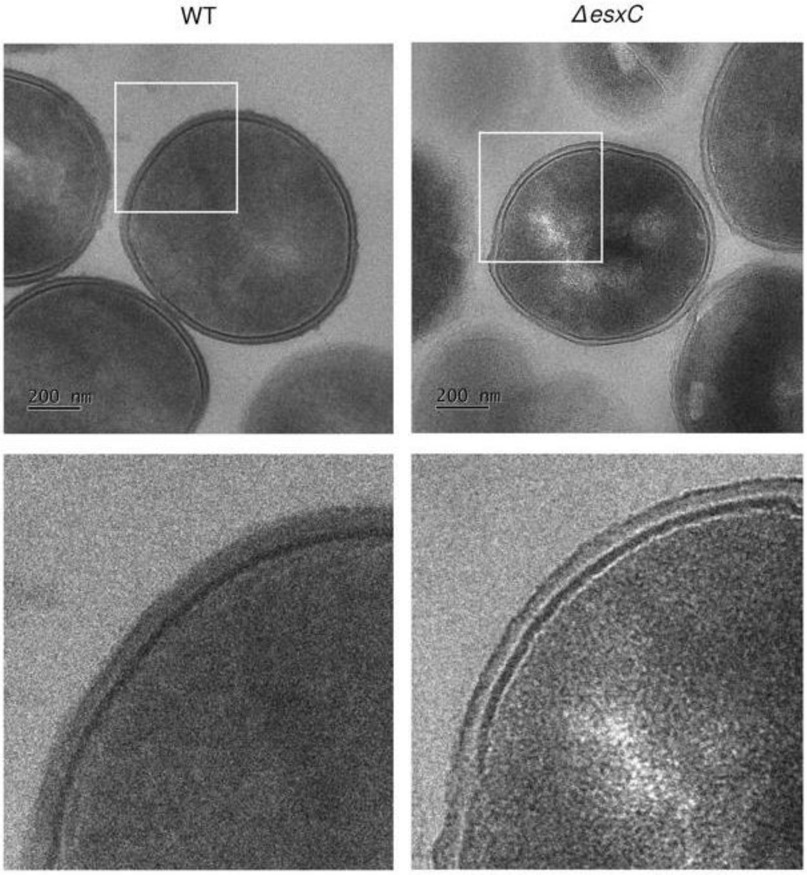


**Figure S4.** Transmission electron micrographs of *S. aureus* USA300 JE2 WT and Δ*esxC* grown to early logarithmic phase

Fig S5

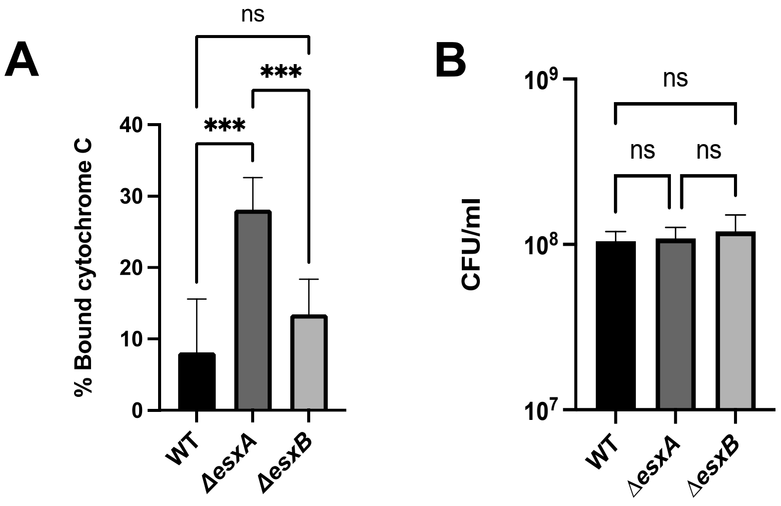


**Figure S5** Quantitative binding assay of cytochrome C to USA300 LAC WT, Δ*esxA* and Δ*esxB* (**A**). The CFU of the starting inoculum was calculated for each mutant **(B)**. Mean ± SD shown, N = 3 (biological replicates), ***P < 0.001 using a one-way ANOVA

### Figure S6

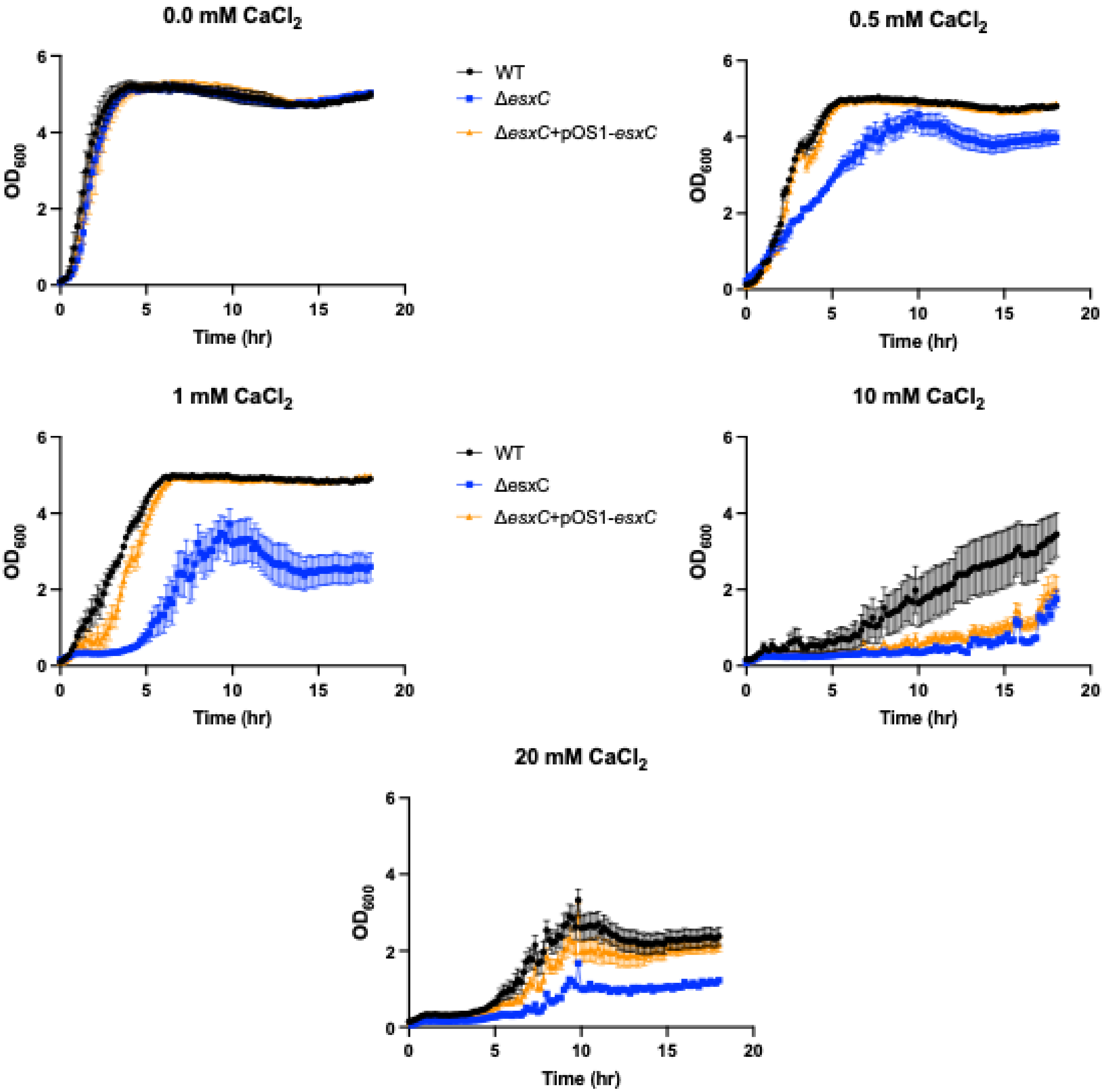


**Figure S6.** Growth curves of WT, *ΔesxC*, *ΔesxC* pOS1-esxC in TSB supplemented with increasing calcium chloride concentrations (0 to 20 mM) in the presence of 5 µg/ml daptomycin. Mean +SE is shown

Fig S7

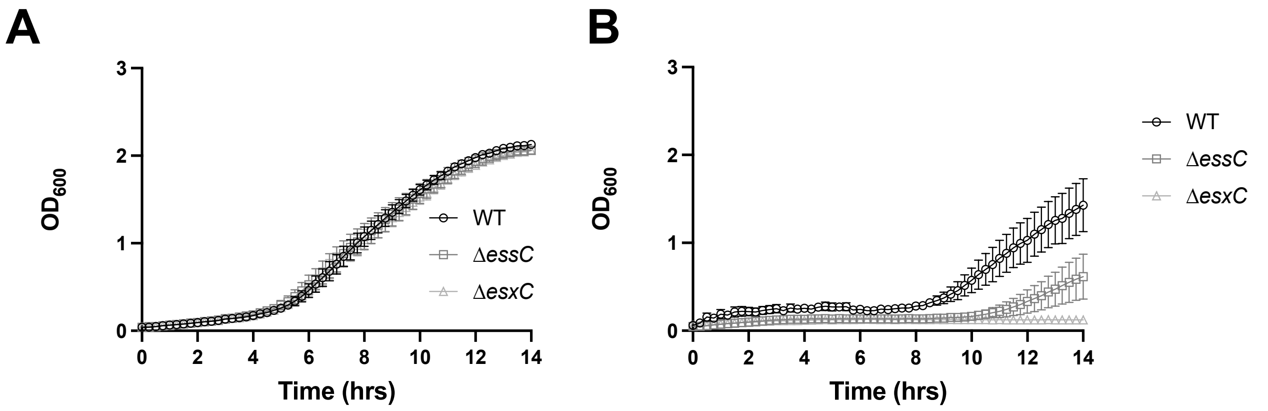


**Figure S7.** (A) Growth curves of WT, Δ*essC* and Δ*esxC* in DMEM + FBS. **(B)** Growth curves of WT, Δ*essC* and Δ*esxC* in DMEM + FBS in the presence of 1µg/ml daptomycin and 1 mM $CaCl_2$. Graphs show mean and SEM of three independent experiments.

Fig S8

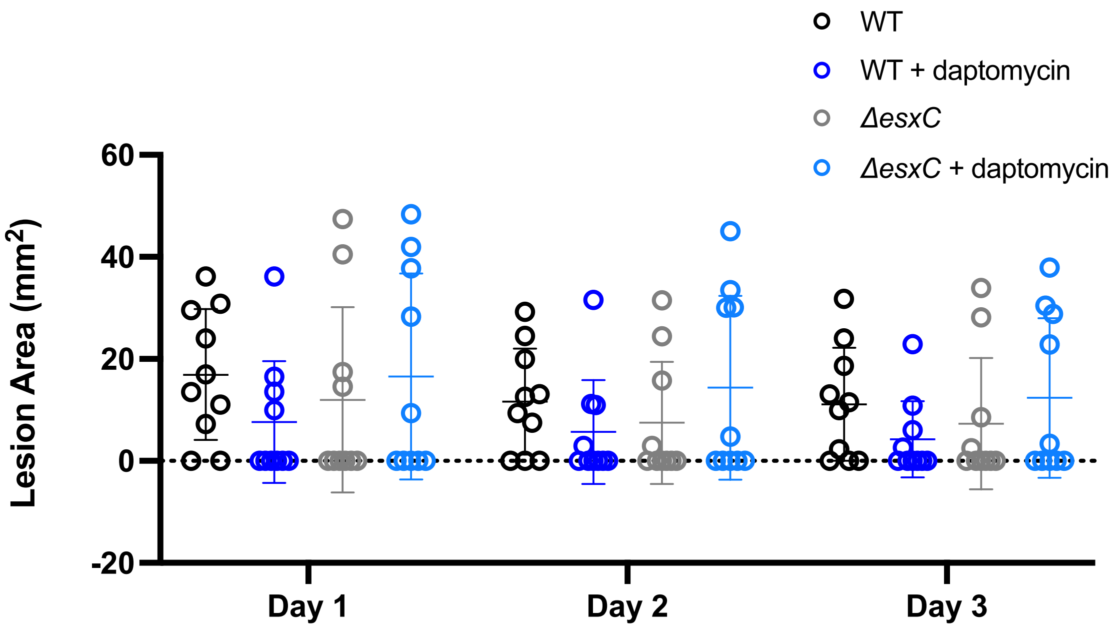


**Figure S8** The area of lesions was measured and calculated daily from mice infected with *S. aureus* WT or esxC mutants. Area of lesions was measured and calculated daily. Graph shows mean ± SD (n = 5, 2 lesions/animal). Two-way ANOVA between groups revealed no significant differences between treatments.
